# mRNA trans-splicing dual AAV vectors for (epi)genome editing and gene therapy

**DOI:** 10.1101/2023.02.07.527504

**Authors:** Lisa Maria Riedmayr, Klara Sonnie Hinrichsmeyer, Stefan Bernhard Thalhammer, Nina Karguth, Sybille Böhm, David Manuel Mittas, Valentin Johannes Weber, Dina Otify, Victoria Splith, Manuela Brümmer, Nanda Boon, Jan Wijnholds, Verena Mehlfeld, Stylianos Michalakis, Stefanie Fenske, Martin Biel, Elvir Becirovic

## Abstract

Large genes including several CRISPR-Cas modules, such as gene activators (CRISPRa), require dual adeno-associated viral (AAV) vectors for an efficient *in vivo* delivery and expression. Current dual AAV vector approaches have important limitations, e.g., low reconstitution efficiency, production of alien proteins, or low flexibility in split site selection. Here, we present a dual AAV vector technology based on <u>re</u>constitution <u>v</u>ia m<u>R</u>NA trans-splicing (REVeRT). REVeRT is flexible in split site selection and can efficiently reconstitute different split genes in numerous *in vitro* models, in human organoids, and *in vivo*. Furthermore, REVeRT can functionally reconstitute a CRISPRa module targeting genes in various mouse tissues and organs in single or multiplexed approaches upon different routes of administration. Finally, intravitreal supplementation of *ABCA4* via REVeRT improves retinal degeneration and function in a mouse model of Stargardt disease. Due to its flexibility and efficiency REVeRT harbors great potential for basic research and clinical applications.

## Introduction

Adeno-associated virus (AAV)-mediated gene transfer is the most efficient and widely used method for *in vivo* gene delivery in mammals^1, 2^. Despite numerous advantages over other therapeutic vectors (e.g., low overall pathogenicity and toxicity and ability to transduce non-dividing cells), AAVs have an important limitation: their low genome packaging capacity of 4.7-5.0 kb^3^. This hampers their use for expression of large genes including many therapeutically relevant CRISPR-Cas9 modules, such as gene activators (CRISPRa), gene inhibitors (CRISPRi), and base or prime editors^2, 4^. To overcome this constraint, several strategies have been established following the same principle: The gene of interest is split into two parts and packaged into separate AAV vectors, typically referred to as dual AAVs. Depending on the approach, the split parts of the gene can subsequently be reconstituted at the AAV genome, mRNA, or protein level upon co-delivery of the dual AAVs to the target tissue^2, 5^.

Gene reconstitution at the AAV genome level (from here on referred to as DNA trans-splicing) was the first dual AAV vector approach applied *in vivo*. The different versions of this method rely on homologous recombination and/or concatemerization of the AAV genome^6–9^. However, DNA trans-splicing often results in low to moderate reconstitution efficiency^10–13^. By comparison, the split intein approach operates at the protein level and, if successful, typically leads to high reconstitution efficiencies^11, 12, 14–17^. Nevertheless, as it requires proper protein folding of both split fragments, this method strongly depends on the split position and relies on the presence of specific amino acids within the split site^18, 19^. Therefore, finding a suitable and efficient split position requires an elaborate and time-consuming screening process for each gene and might fail completely. In addition to its low flexibility, this approach creates equimolar amounts of potentially immunogenic or pathogenic proteins (inteins), raising important safety concerns for translational approaches. This may become even more relevant if split inteins are combined with Cas-based technologies, given that a co-expression could enhance host immune response in an additive or synergistic manner.

Consequently, there is a high unmet need for a strategy that overcomes the limitations of the current gene reconstitution approaches. Reconstitution via dual mRNA trans-splicing AAVs remains a largely unexplored option. Despite utilizing similar sequence elements as DNA trans-splicing (e.g., complementary domains and splice sites), the reconstitution in the mRNA trans-splicing approach occurs between two different mRNA molecules, i.e., in *trans*^20^. Thus, it is fundamentally different from DNA trans-splicing, in which mRNA splicing occurs in *cis* with the sole purpose to remove the non-coding portions between the split fragments after successful reconstitution at the genome level. Nevertheless, in many studies, dual DNA trans-splicing AAV vectors have been incompletely termed trans-splicing^21–23^ or were even mixed up with mRNA trans-splicing approaches^11^. This unfavorable terminology makes it difficult for non-experts to distinguish between the respective methods.

Here, we develop flexible and efficient dual AAV vectors utilizing mRNA trans-splicing for reconstitution of large coding sequences. These vectors have been tested in the context of various therapeutically highly relevant genes or gene editing modules in a variety of challenging experimental setups *in vitro* and *in vivo*.

## Results

### Development of REVeRT technology for reporter gene reconstitution in vitro

To test and optimize REVeRT, we developed a split-fluorophore assay that results in expression of cerulean, a cyan fluorescent protein, upon successful reconstitution of the two split fragments (Fig. 1a). For this experiment, we used the CAG|GT sequence as a split site for REVeRT with the vertical line symbolizing the split point. This sequence contains the exonic parts of splice site sequences, i.e., CAG for the splice donor site followed by G for the splice acceptor site. The T at the last position is part of the exonic sequence of a very strong splice acceptor published in our recent study^24^. The CAGGT sequence cannot be found within the short (720 bp) native cerulean coding sequence. However, this sequence can be easily created by using synonymous codons without changing the protein code (Fig. S1a, split site #1).

**Fig. 1.**
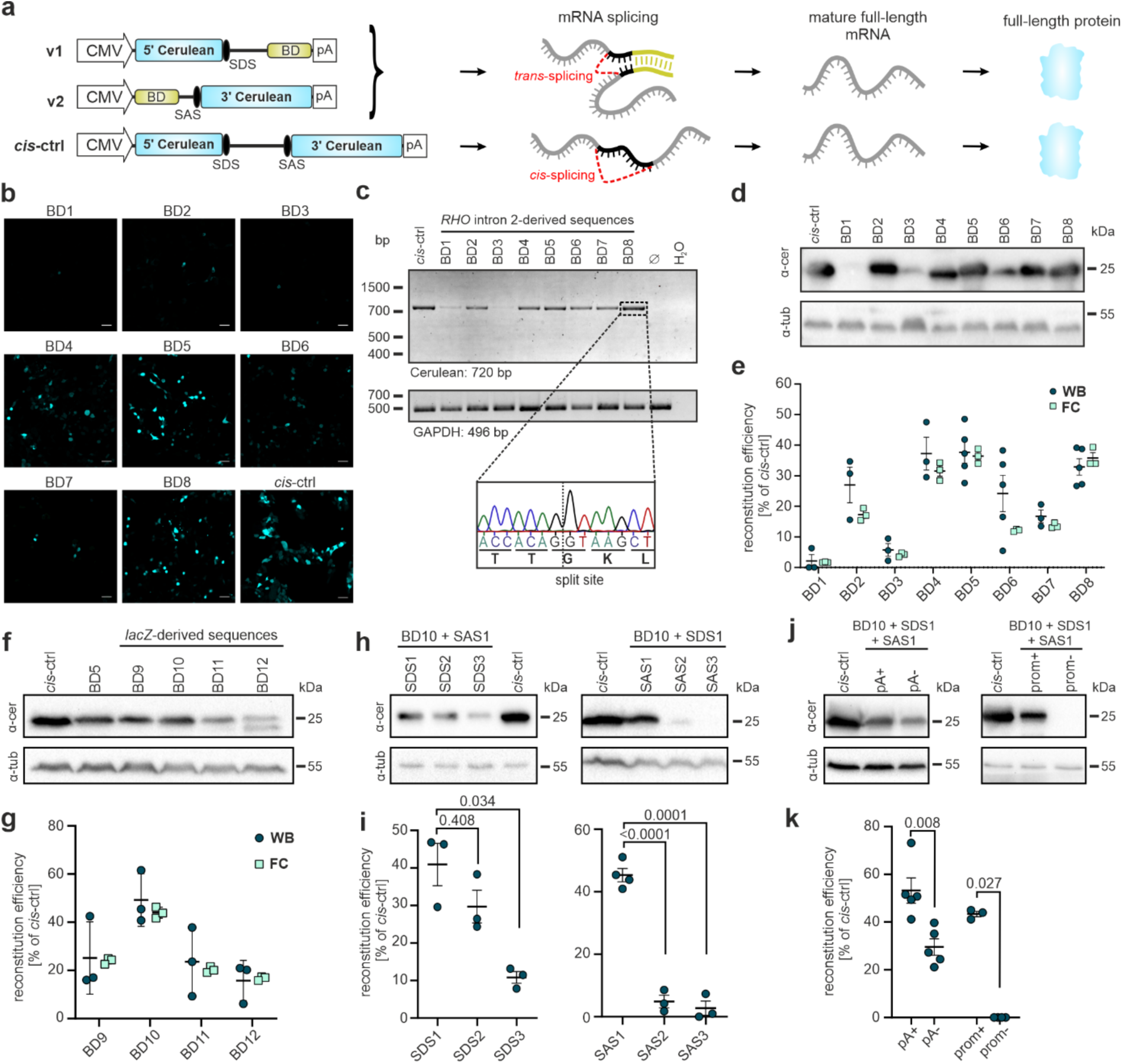
Development and optimization of mRNA trans-splicing *in vitro*. **a** Illustration of split fluorophore assay to test reconstitution via mRNA trans-splicing. A cis-splicing vector was used as a positive control (cis-ctrl). v1, vector 1; v2, vector 2; BD, binding domain; SDS, splice donor site; SAS, splice acceptor site; pA, polyadenylation signal. **b** Confocal images of HEK293 cells transfected with v1 and v2 containing different BDs. Scale bar: 50 µm. **c** RT-PCR of reconstituted full-length cerulean. GAPDH serves as input reference. Sequencing result of the splicing junction is shown below. bp, base pairs. **d** Western blots of transfected cells shown in **b**. α-cer, anti-cerulean antibody; α-tub, anti-β-tubulin antibody; kDa, kilodalton. **e** Quantification of reconstitution efficiency relative to cis-ctrl using western blots (WB, dark blue) or flow cytometry (FC, light blue). **f** Western blot of cells transfected with v1 and v2 containing different human (*RHO* intron 2-derived, BD 5) or bacterial (*lacZ*-derived, BD 9-12) binding domains. **g** Ratiometric quantification of reconstitution efficiency of the transfections shown in **f** relative to cis-ctrl. n = 3. **h** Western blots of cells transfected with v1 and v2 containing different SDS or SAS. **i** Ratiometric quantification of reconstitution efficiencies relative to cis-ctrl. Welch ANOVA with Dunnett T3 (SAS) or Kruskal-Wallis test with Dunnett T3 (SDS) was used. **j** Western blots of cells transfected with versions of v1 containing (+pA) or lacking (-pA) a polyadenylation signal and with versions of v2 containing (+prom) or lacking (-prom) a promoter. **k** Ratiometric quantification of reconstitution efficiencies relative to cis-ctrl. n = 3-5. Mann-Whitney test was used. Scatter plots show mean ± SEM.

Both split fragments were equipped with elements required for transcript expression and subsequent mRNA trans-splicing, i.e., promoter, complementary binding domains, splice sites and polyadenylation signal. For initial testing, we used binding domain sequences of different lengths derived from the intronic sequence of the human rhodopsin (*RHO*) gene (Fig. S1b). Native intronic sequences typically contain intronic splicing enhancers that may facilitate mRNA trans-splicing-mediated reconstitution. We examined the principal reconstitution capability of mRNA trans-splicing in transfected HEK293 cells and found that it results in a seamless, precise and efficient ligation of the two fragments (Fig. 1b-e). Nevertheless, although helpful at this initial stage, human-derived sequences as binding domains are less suitable for translational approaches, as they are expected to bind to the respective human transcripts increasing the risk of off-target effects. Therefore, we tested additional binding domains derived from a 100 bp sequence of the bacterial *lacZ* gene that was modified to create four distinct binding domain sequences without any homology to the human or mouse genome. One of them resulted in the highest reconstitution efficiency observed (44.0 % in flow cytometry and 53.5 % in western blot-based quantifications, Fig. 1f, g and Table S1) and was therefore considered as the optimized binding domain for further experiments.

Next, we analyzed other trans-splicing elements for their effect on the reconstitution efficiency, i.e., position of the split site, presence of the polyadenylation signal, and strength of splice sites (Fig. 1h-k and Fig. S1c, d). An element showing a high impact on the reconstitution efficiency was the splice acceptor site (SAS). When comparing three different SASs predicted to be strong using splice site prediction software, efficient reconstitution was only observed for one SAS variant (Fig. 1h, i). This SAS was recently shown to be highly effective in different conditions^24^. Compared to the SAS, the splice donor site (SDS) showed a less pronounced effect as well as the presence of a polyadenylation signal (Fig. 1j, k). No cerulean reconstitution was observed when the promoter was removed from the 3’ split fragment (v2) (Fig. 1j, k). This suggests that reconstitution is driven exclusively by mRNA trans-splicing and not via processes which occur at the DNA level such as homologous recombination. In another set of experiments, we show that different split sites lacking a C at the first and/or T at the last position of the CAGGT sequence all lead to a successful reconstitution yielding moderate to high expression levels (Figure S1c, d). This indicates that AGG is the minimal split site consensus sequence for REVeRT. Moreover, REVeRT can be used for a directional reconstitution of the fluorophore sequence split into three fragments facilitated by two different binding domains (Fig. S1e, f) and is independent of the cell type (Fig. S1g, h).

Conclusively, in this part of the study, we identified and optimized several molecular determinants that affect the efficiency of REVeRT. These optimized elements (Table S2) were used for all subsequent experiments.

### Evaluation of dual REVeRT AAVs in vivo

Next, we analyzed the performance of REVeRT in a dual AAV vector setting *in vivo*. The retina is the target organ of one of the first approved AAV gene therapies and for many others currently in clinical development^25, 26^. However, more than 30 inherited retinal disorders are caused by mutations in large genes exceeding the AAV vector DNA capacity (https://sph.uth.edu/retnet/). We therefore evaluated the functionality of REVeRT in retinal cells by injecting two different split fluorophore variants into the eyes of C57BL/6J wild-type (WT) mice using AAV8(Y733F) vectors (Fig. S2a, b)^27^. The first variant expressing the split cerulean cassette was injected subretinally to target photoreceptors. The second variant contains one additional fluorophore, citrine or mCherry, added to each of the split fragments to monitor the expression of the individual vectors and was injected under the monolayer of retinal pigment epithelial (RPE) cells. Four weeks after injection, both approaches showed robust and specific fluorophore expression in the respective target cells (Fig. S2c-e).

Next, to monitor REVeRT-mediated reconstitution in different organs, we developed a split luciferase reporter system. Three different split sites in the luciferase sequence were tested *in vitro* (Fig. S3). In line with the split cerulean assay, the different split sites showed only minor differences in reconstitution efficiency, further underlining the high flexibility of REVeRT in the split site selection. No luciferase signal was obtained for single split vectors (Fig. S3b). We used the REVeRT luciferase system to evaluate the reconstitution efficiency in different organs upon intraperitoneal application. These are very challenging conditions considering that AAVs are diluted in the bloodstream, have to cross several biological barriers, and co-transduce the same cells before mRNA trans-splicing can occur. Three weeks after injecting dual AAV9 vectors into WT mice, luciferase signal was detectable throughout the body (Fig. 2a, b). The reconstituted luciferase was expressed in various organs of the injected animals with the strongest signal detectable in the heart (Fig. 2c).

**Fig. 2.**
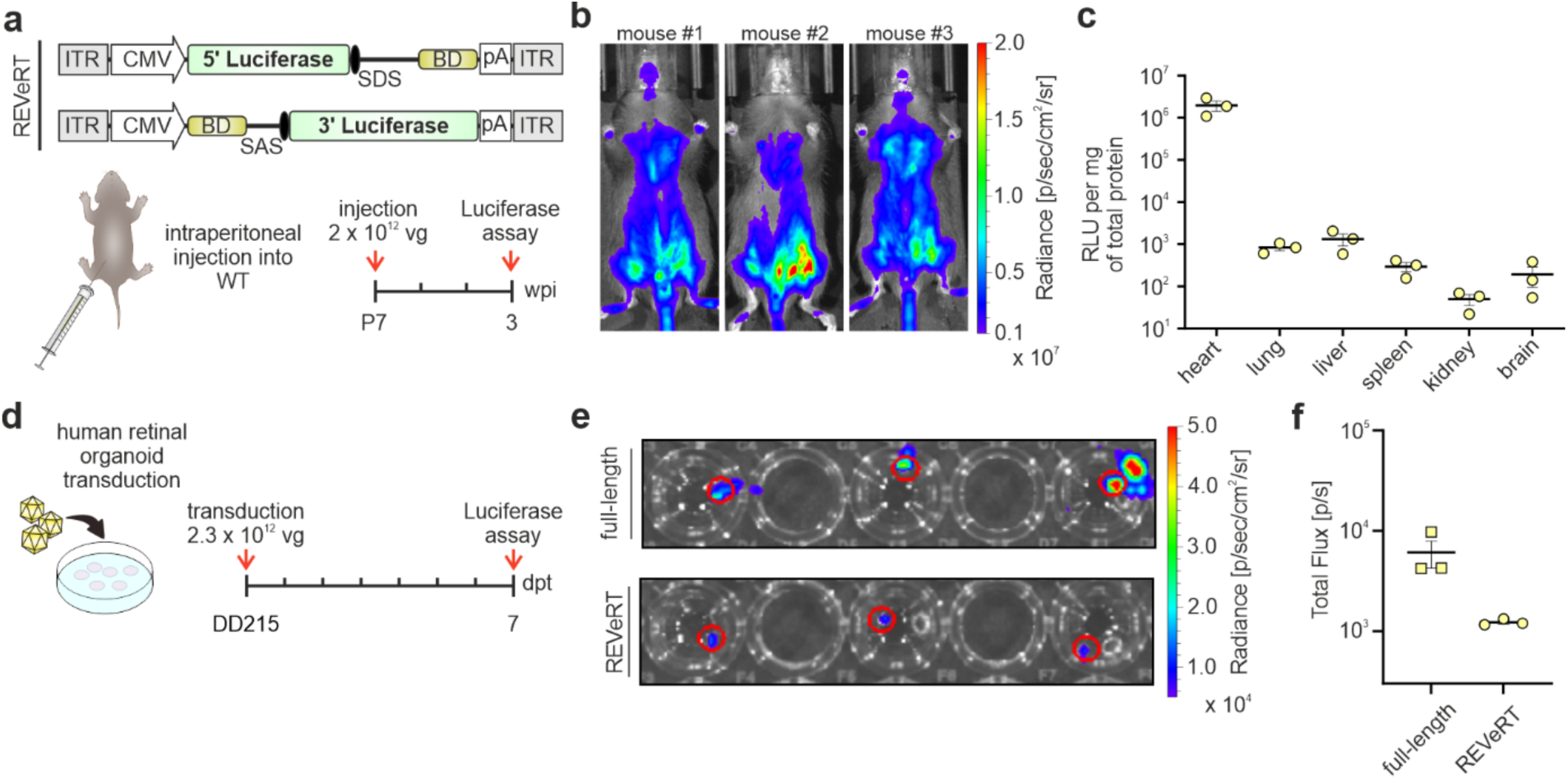
Reconstitution of luciferase via dual REVeRT AAVs *in vivo* and in human retinal organoids. **a** Dual REVeRT AAVs expressing a split luciferase gene. Lower panel, Time scale for experiments in WT mice (**b**, **c**). **b** Luminescence obtained from WT mice co-injected with titer-matched dual REVeRT AAV9 vectors. **c** Relative quantification of luciferase expression in different organs three weeks post-injection. RLU, relative light units. **d** Time scale for experiments in human retinal organoids transduced with dual REVeRT AAV9 vectors at differentiation day 215 (DD215). dpt, days post transduction. **e** Luminescence obtained from transduced organoids. Regions of interest indicating the organoid position were used for quantification and are shown in red. A AAV vector expressing full-length luciferase served as positive control. **f** Quantification of the results shown in **e**. Scatter plots show mean ± SEM. n = 3 mice or retinal organoids.

Collectively, these results show that dual REVeRT AAVs lead to the reconstitution of several split reporter genes in mice after local or systemic application of different AAV capsids.

### Evaluation of dual REVeRT AAVs in human retinal organoids

To better assess the translational potential of dual REVeRT AAVs, we used human retinal organoids, which are frequently employed to evaluate the potency of therapeutic AAV vectors. To this end, we co-transduced human retinal organoids with AAV9 vectors carrying the split luciferase gene (Fig. 2d). The reconstituted luciferase reached 20.2 % of the signal from control organoids transduced with single AAVs expressing the full-length luciferase (Fig. 2e, f). This demonstrates that dual REVeRT AAVs can be used to transduce highly differentiated human tissue cultures.

### Dual REVeRT AAVs result in functional reconstitution of dCas9-VPR in vitro and in vivo

Having studied the efficacy of REVeRT using reporter assays, we set out to evaluate this technology in a therapeutically relevant context. For this purpose, we focused on “dead” (d)Cas9-VPR, a very potent CRISPRa module with high translational potential^28^. The range of applications of CRISPRa for the treatment of genetic or acquired diseases includes transcriptional activation (transactivation) of silent genes that are functionally equivalent to defective genes^29^. The *MYO7A* (*USH1B*) gene represents an attractive target for CRISPRa therapy, as it exceeds the capacity of AAV vectors and is associated with Usher syndrome (USH), the most common form of genetic deafblindness. *MYO7A* possesses a functionally equivalent counterpart, *MYO7B*, in humans and in mice fulfilling all requirements for a replacement of *MYO7A*^30, 31^. Following this idea, we first tested single guide (sg)RNAs targeting the region upstream of the *Myo7b* transcriptional start site in 661W cells, immortalized derivatives of mouse cone photoreceptors, co-transfected with split dCas9-VPR. We identified a combination of sgRNAs that showed efficient activation of *Myo7b* (Fig. S4a-d). To analyze whether *Myo7b* can be activated by dual REVeRT AAVs expressing split dCas9-VPR, we co-transduced primary mouse hippocampal neurons with dual AAV8(Y733F) vectors. This resulted in robust dCas9-VPR reconstitution accompanied by strong *Myo7b* expression (Fig. S4e, f).

The promising *in vitro* results encouraged us to test whether dual REVeRT AAV vectors are capable of functionally reconstituting dCas9-VPR *in vivo* under more challenging conditions. As a read-out for dCas9-VPR function, we analyzed the activation of *Myo7b* using the sgRNAs identified in 661W cells. For this purpose, adult WT mice were subretinally injected with dual REVeRT AAV8(Y733F) vectors (Fig. 3a, b). Four weeks after injection, we observed a strong activation of *Myo7b* at transcript and protein level (Fig. 3c-e). Next, we tested the same AAV vectors in other organs using different routes of administration. A stereotactic injection into the hippocampus of adult WT mice also resulted in robust activation of the *Myo7b* gene four weeks after injection (Fig. 3f, g). In both retina and hippocampus, the *Myo7b* expression correlated with the dCas9-VPR reconstitution efficiency, suggesting a dose dependency of the transactivation (Fig. S4g, h). We also observed efficient activation of *Myo7b* in several organs (retina, heart, liver, skeletal muscle, and lung) after intraperitoneal injection of these vectors (Fig. 3h, i). Notably, the highest gene activation was detectable in the heart. This is in line with our results from the luciferase assay, which shows the highest REVeRT-mediated luminescence in this organ.

**Fig. 3.**
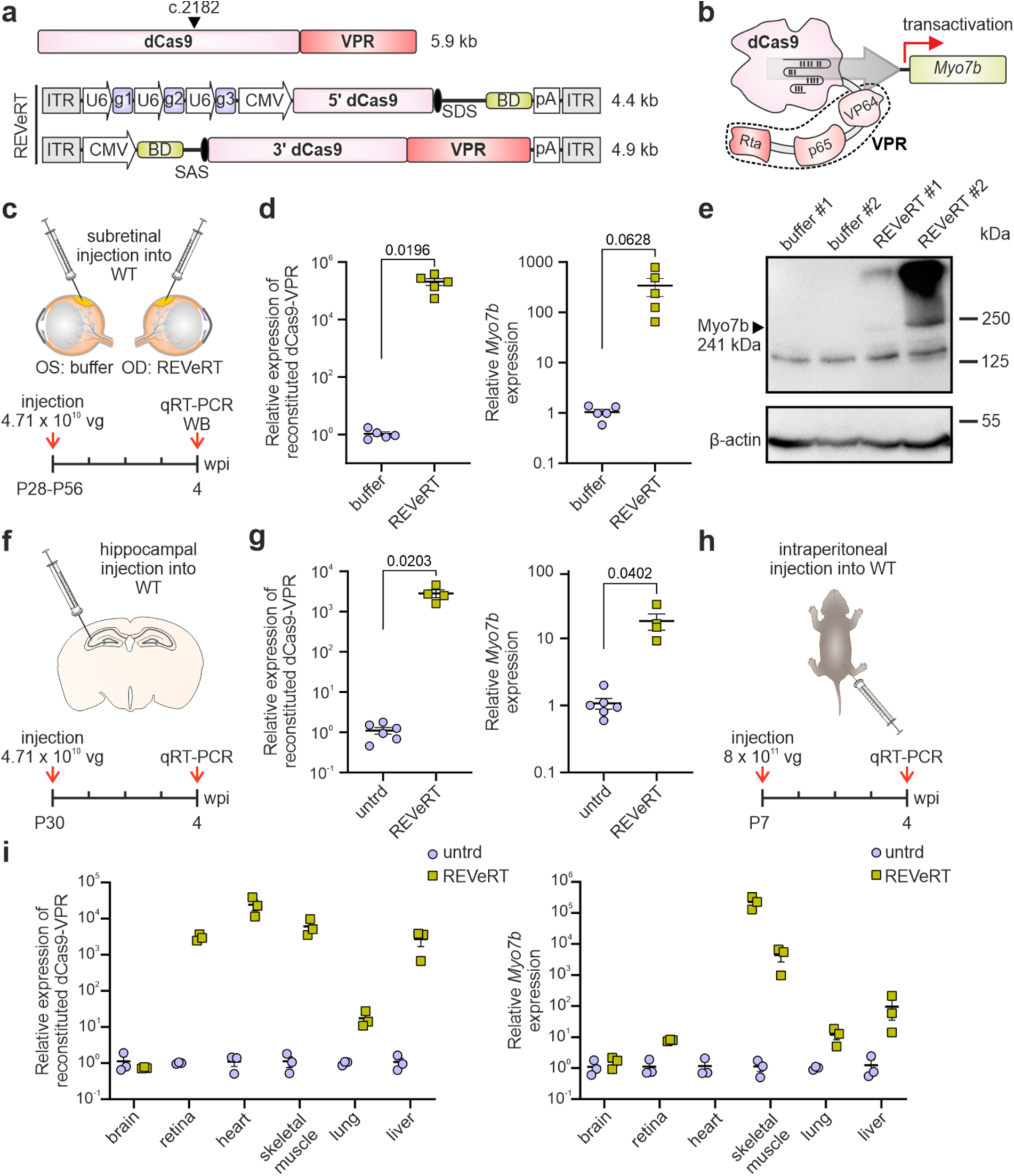
Reconstitution of dCas9-VPR in various tissues *in vivo* upon local and systemic application. **a** Upper panel, dCas9-VPR size and the split site used for REVeRT. Lower panel, REVeRT vector cassettes expressing split dCas9-VPR and three sgRNAs (g1 – g3) targeting the promoter region of *Myo7b*. Titer-matched dual REVeRT AAV8(Y733F) vectors were injected via different routes of administration shown in **c**, **f**, and **h**. **b** dCas9-VPR targeting the *Myo7b* promoter. **c** Experimental design for **d**, **e**. OD, *Oculus dexter*; OS, *Oculus sinister*. Contralateral eyes injected with AAV formulation buffer served as reference. **d** qRT-PCR from injected retinas. Shown are the relative expression levels of reconstituted dCas9-VPR (left) and the transactivated *Myo7b* (right). **e** Western blot from the retinas of two mice (#1 and #2) injected with REVeRT AAVs or buffer. **f** Experimental design for **g**. **g** qRT-PCR from injected hippocampi. Shown are the relative expression levels of reconstituted dCas9-VPR (left) and transactivated *Myo7b* (right). Untreated (untrd) hippocampi served as reference. **h** Experimental design for **i**. **i** qRT-PCR from organs or tissues harvested of intraperitoneally injected mice. Shown are the relative expression levels of reconstituted dCas9-VPR (left) and transactivated *Myo7b* (right). Untrd organs served as reference. A two-tailed unpaired t test with Welch’s correction was used. Scatter plots show mean ± SEM. n = 3-5 organs/tissues.

Taken together, these results demonstrate that the REVeRT technology results in functional reconstitution of dCas9-VPR *in vivo*, irrespective of the administration route.

### REVeRT leads to functional reconstitution of prime editors in vitro

In addition to CRISPRa, prime editors (PEs) represent another CRISPR-Cas module that exceeds the packaging capacity of AAVs (Fig. S5a) and has high translational potential^32, 33^. With the goal of developing dual REVeRT AAV vectors that can reconstitute PEs *in vivo*, we first attempted to edit the mouse *Dnmt1* locus using previously published prime editing guide (peg) RNAs^32^. Nevertheless, even when expressing the full-length prime editor in transfected 661W cells, no editing of this locus was detectable (Fig. S5b). Similarly, in our attempts to modify other mouse genes (*Rho*, *Cnga1* or *Ush2a*) under different conditions, e.g., in presence of full-length PE2 and PE3^32^, different pegRNAs, or different murine cell lines, we did not observe any editing of these loci (Fig. S5b). This suggest that mouse loci may be less susceptible to prime editing and prompted us to analyze prime editing of the intergenic human *HEK3* locus using published pegRNAs designed to insert CTT^32^. When the full-length PE2 was expressed in HEK293T cells, we observed an editing efficiency of 18 %, which is comparable to previously reported results^32^. Next, we co-expressed dual REVeRT PE2 in HEK293T cells. The resulting editing efficiency (16 %) was similar to that of full-length PE2 (Fig. S5c). This demonstrates that REVeRT is able to functionally reconstitute another therapeutically relevant CRISPR-Cas module *in vitro*.

### In vivo multiplexing for simultaneous gene knockout and activation using dual REVeRT AAVs

It has recently been shown that catalytically active Cas9-VPR can be used for simultaneous activation and knockout of different genes. This is based on the finding that Cas9 loses its catalytic activity in combination with short (< 16 nt) sgRNA spacers^34, 35^. Consequently, **con**current k**n**ockout and **act**ivation (hereafter referred to as CONNACT) of different genes can be achieved by combining sgRNAs of different spacer lengths targeting the respective loci. Here, we investigated whether REVeRT is suitable for such multiplexing approaches. One application for CONNACT is the treatment of gain-of-function mutations, which require an efficient knockdown of the diseased allele and the supplementation of another gene to compensate for the missing function. Gain-of-function mutations are frequently found in the rhodopsin gene (*RHO*), including the most common Pro23His (P23H) substitution, and have been associated with retinitis pigmentosa^36^. Moreover, it has recently been shown that transactivation of the M-opsin-encoding gene (*Opn1mw*) can compensate for the lack of rhodopsin function in a mouse model of retinitis pigmentosa^15^. In an attempt to develop a multiplexing approach targeting such mutations, we used dual REVeRT AAV8(Y733F) vectors expressing split Cas9-VPR to simultaneously knockout *Rho* and activate *Opn1mw*. For this purpose, we first tested a number of different 15 nt spacer sgRNAs targeting the region upstream of the *Opn1mw* transcriptional start site in mouse embryonic fibroblasts (MEFs) (Fig. S5d, e). Notably, in contrast to the results of a recent study targeting other genes^34^, we found that 15 nt spacer sequences consistently lead to a decrease in transactivation efficiency compared to their 20 nt counterparts (Fig. S5f).

Considering the multifaceted readout, we concluded that reconstitution of Cas9-VPR in combination with the CONNACT strategy is most appropriate to compare the efficiency of REVeRT and split-intein-mediated reconstitution *in vivo*. Therefore, adult WT mice were injected with the dual REVeRT or split-intein AAV cassettes containing one sgRNA with a 20 nt spacer (g1) for *Rho* knockout and the two sgRNAs with a 15 nt spacer (g2+g3) for *Opn1mw* transactivation (Fig. 4a, b). Four weeks after injection, both, gene knockout and gene activation, were detected in the injected retinas and there were no obvious differences in efficiency when comparing REVeRT and split inteins (Fig. 4c). Next, we evaluated this multiplexing approach in the P23H mouse model of autosomal dominant retinitis pigmentosa by injecting dual REVeRT AAV vectors subretinally into heterozygous *Rho*^P23H/+^ mice. In this context, we also investigated whether the transactivation efficiency could be further increased by adding another *Opn1mw*-targeting sgRNA (g4) to the expression cassette. Four weeks post injection, both, *Rho* knockdown and *Opn1mw* transactivation, were detected at the transcript and protein level (Fig. 4d, e). The introduction of a third sgRNA targeting *Opn1mw* resulted in a slight increase in activation efficiency (Fig. 4d). RNA-Seq of the injected retinas confirmed the *Opn1mw* activation and *Rho* knockdown (Fig. S5g) and revealed that only a small fraction of additional genes was up-or downregulated under these conditions (Table S3).

**Fig. 4.**
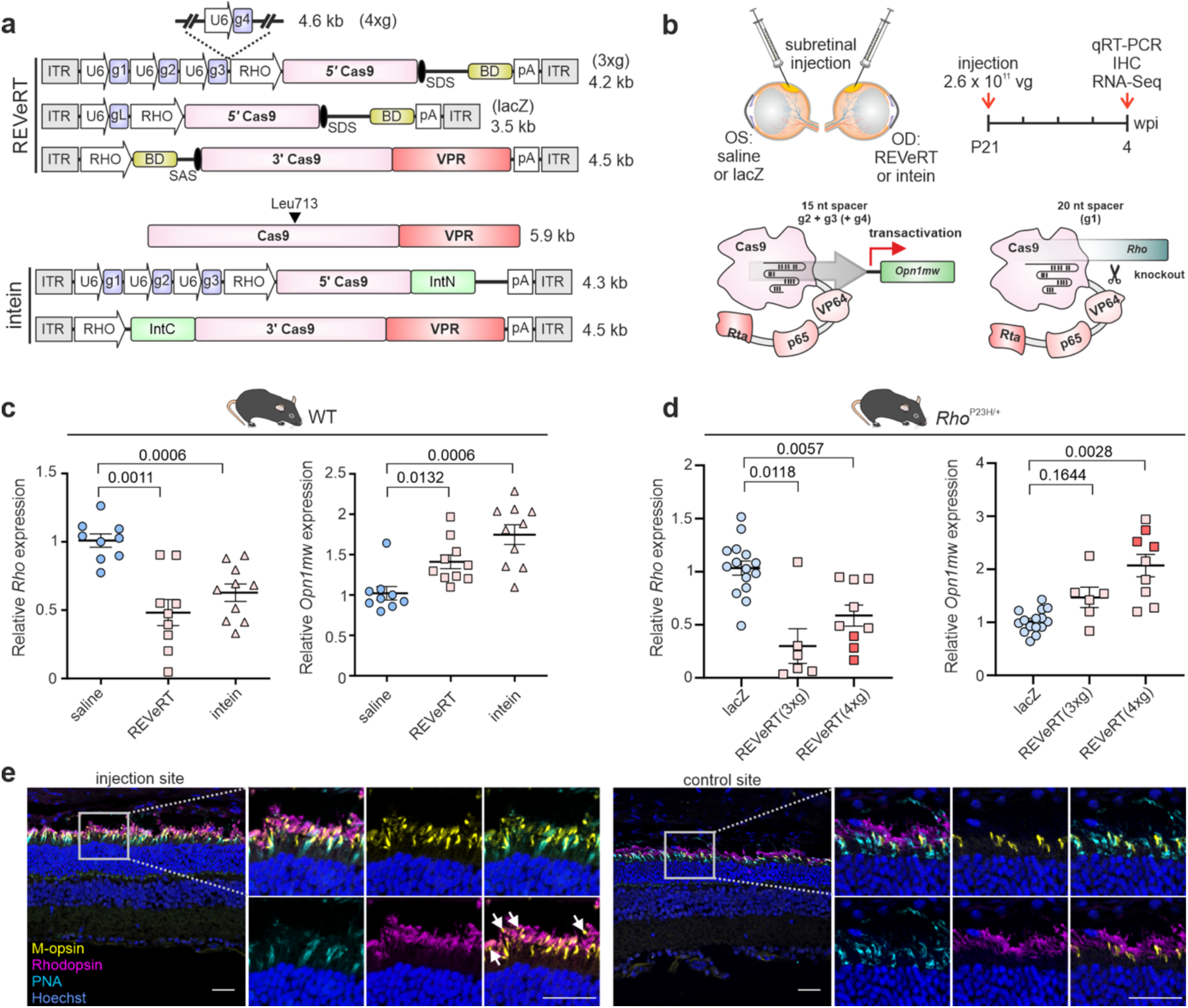
Simultaneous gene knockout and transactivation using REVeRT. **a** Upper panel, Dual REVeRT AAV8(Y733F) vector cassettes expressing split Cas9-VPR and three sgRNAs targeting either exon 1 of *Rho* (g1) or the promoter region of *Opn1mw* (g2-g4). Lower panel, The corresponding dual split intein AAV8(Y733F) vector cassettes used for side-by-side comparison to REVeRT shown in **c**. **b** Experimental design and timeline of experiments shown in **c**-**e**. sgRNAs with different spacer lengths are used for a simultaneous knockout (20 nt, g1) of *Rho* and transcriptional activation (15 nt, g2-g4) of *Opn1mw*. The fourth sgRNA (g4) was added in **d** and **f** as indicated. **c** Relative expression of *Rho* and *Opn1mw* in retinas of injected WT mice. Retinas injected with saline solution were used as a reference. n = 9-10 mice. **d** Relative expression of *Rho* and *Opn1mw* upon injection of *Rho*^P23H/+^ mice with one of two versions of dual REVeRT vectors expressing g1 and either two (REVeRT(3xg)) or three (REVeRT(4xg)) *Opn1mw* transactivating sgRNAs. lacZ-injected retinas were used as a reference. The three retinas used for RNA-Seq analysis (Fig. S5g) are shown in dark pink. Welch ANOVA with Dunnett T3 was used for statistical analysis in **c** and **d**. n = 6-9 mice. **e** Immunostainings of retinal cryosections of *Rho*^P23H/+^ mice subretinally injected with REVeRT(3xg) (left). A control site distal from the injection site is shown as a reference (right). Magnification of areas marked by white rectangle are shown. White arrowheads indicate rod outer segments expressing M-opsin while lacking any rhodopsin signal. Peanut agglutinin lectin (PNA) was used as a cone outer segment marker. In the control site, M-Opsin is only expressed in cones. Scale bar: 30 µm. Scatter plots show mean ± SEM.

In summary, these data demonstrate that dual REVeRT can be used for CRISPR-Cas-based multiplexing approaches designed to treat complex diseases requiring simultaneous activation and downregulation of different genes.

### Gene supplementation therapy for inherited blindness using dual REVeRT AAVs

Finally, we evaluated the potency of dual REVeRT AAV vectors in the context of gene supplementation therapy. For this, we focused on the treatment of the Abca4-deficient (*Abca4*^-/-^) mouse model for Stargardt disease, which represents the most common inherited macular degeneration^37^. To date, no therapy for ABCA4-related Stargardt disease exists, although both split-intein and DNA trans-splicing have been evaluated for their therapeutic potency in mouse models and pigs^12, 13, 38, 39^. The *Abca4*^-/-^ mouse model shows only a mild retinal phenotype^40^, which complicates the efficacy evaluation of gene therapies. However, disease progression is enhanced in mice lacking Rdh8 in addition to Abca4^41, 42^. To determine the optimal injection time point, we first characterized the retinal degeneration in this *Abca4^-/-^Rdh8^-/-^* mouse model over a period of three months. Using optical coherence tomography (OCT) and electroretinography (ERG), we detected the strongest retinal degeneration between postnatal week (PW) 3 and 5 (Fig. S6a, b). Animals were injected at PW2 with dual REVeRT AAVs expressing split *ABCA4* (Fig. 5a, b). We chose an intravitreal injection using the recently described AAV2.GL capsids^43^. Although intravitreal injections are a more challenging route of administration as AAVs must overcome additional barriers, they are therapeutically attractive as they are less invasive and require less technical skill or specialized equipment^44^ than subretinal injections. Four weeks post injection, no differences could be seen between the injected and control eyes (Fig. S6c-f). Ten weeks after injection, however, we observed improvement in retinal structure and function in three out of four treated eyes compared to the contralateral control eyes injected with the AAV formulation buffer (Fig. 5c, d; Fig. S7a-d). In three animals (mouse #1, #2 and #3), these improvements were pronounced, suggesting that REVeRT can lead to fully functional *ABCA4* reconstitution even after intravitreal injection (Fig. 5d). Another hallmark for retinal degeneration in *Abca4*^-/-^ mice and in patients is the accumulation of bisretinoid-containing lipofuscin pigments in the RPE, which can be occasionally detected as distinct white dots in scanning laser ophthalmology (SLO) images^45, 46^. In mouse #1, which showed the highest improvement in ERG and OCT measurements, we detected such dots in the control eye but not in the treated eye (Fig. 5e). However, SLO images of both, control and treated eyes, of the remaining mice did not show white dots (Fig. S7e). To investigate whether the observed treatment effects correlate with protein expression, we performed immunostainings or western blots with the injected retinas. Indeed, all mice with improvement in ERGs and/or OCTs showed ABCA4 protein expression, while no signal was detectable in outer segments of buffer-injected contralateral control eyes (Fig. 5f, g). For mouse #4, which showed low to no improvement, no ABCA4 protein expression was detectable by immunostaining (Fig. 5g).

**Fig. 5.**
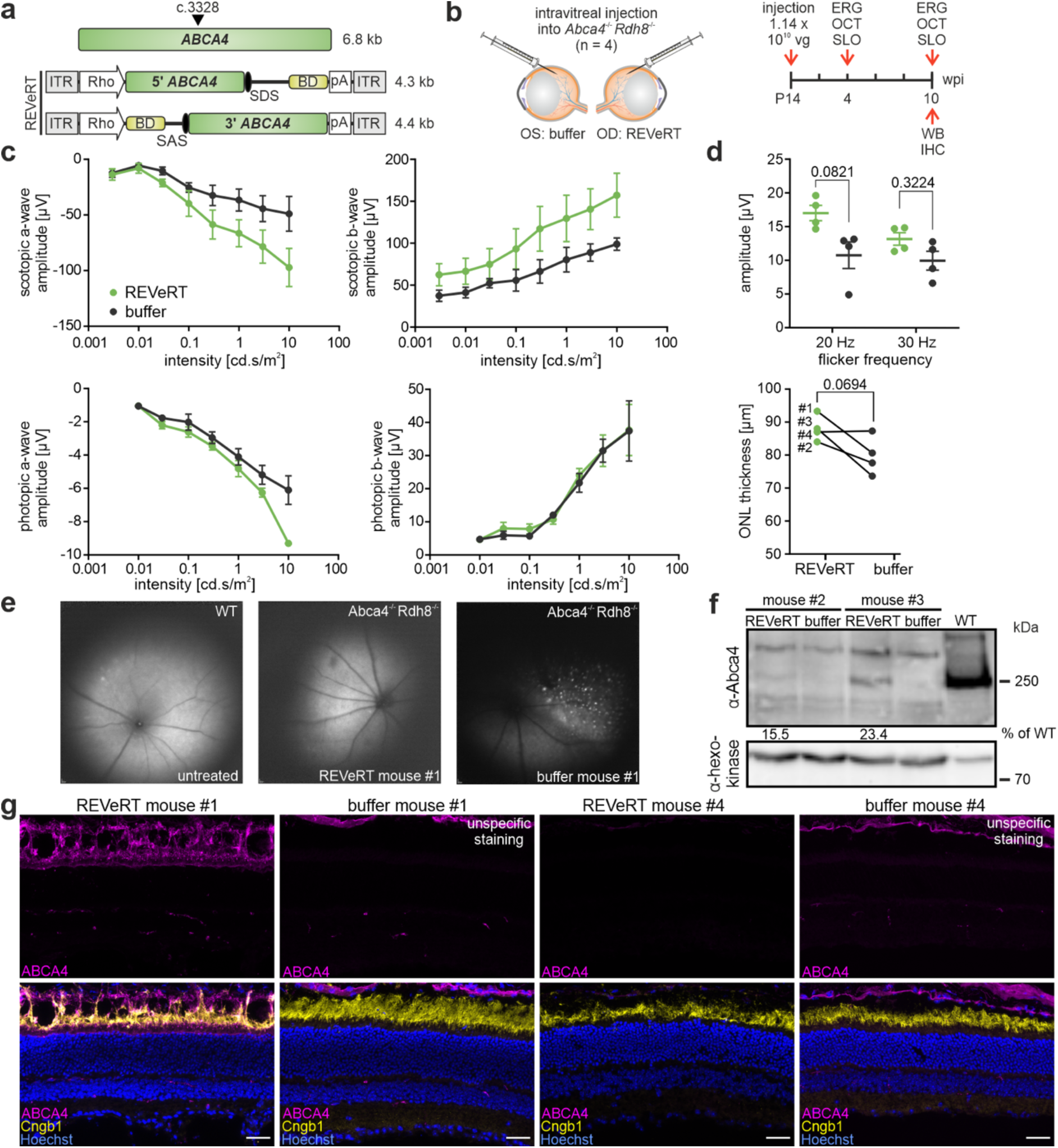
Gene supplementation therapy in a mouse model for an inherited retinal disease using REVeRT. **a** Upper panel, *ABCA4* size and the split site (arrowhead). Lower panel, Dual REVeRT AAV.GL vectors expressing split *ABCA4*. **b** Experimental design for **c**-**g**. **c** ERG measurements under scotopic (upper panel) or photopic (lower panel) conditions at 10 wpi. **d** Upper panel, Results from flicker at 20 Hz and 30 Hz under photopic conditions. Lower panel, Pairwise comparison of OCT measurements the treated and control eye of the single animals. Two-way ANOVA with Šídák’s multiple comparisons test was used for statistical analysis. **e** SLO images from a WT eye and both eyes of mouse #1. **f** Western blot from retinas of mouse #2 and mouse #3 (moderate or strong improvement in OCT and ERG measurements, respectively). **g** Representative immunostainings on retinal sections of mouse #1 (left) and mouse (#4) showing a strong or weak improvement in OCT and ERG measurements, respectively. Upper panel, Images from the right eye injected with dual REVeRT AAVs. Lower panel, Images from the contralateral control eye injected with buffer solution. Scale bar: 30 µm. Cyclic nucleotide-gated channel (Cngb1) antibody was used as a rod outer segment marker to evaluate ABCA4 localization. Some unspecific ABCA4 staining was detected across all sections above the outer segments. Plots show mean ± SEM.

These results demonstrate that the dual REVeRT AAV technology can be used for reconstitution and functional augmentation of a large gene, resulting in therapeutic benefit in a mouse model of inherited blindness.

## Discussion

In this study, we developed a versatile and efficient dual AAV vector technology termed REVeRT that is based on mRNA trans-splicing. We show that REVeRT can be used for functional reconstitution of CRISPRa and prime editors as well as for supplementation of disease-associated genes exceeding the genome packaging capacity of AAVs. In the retina, REVeRT operates at similar efficiencies as split inteins, which are currently the most efficient dual AAV vector approach used in several therapeutically relevant studies^11, 12, 14^. However, REVeRT offers at least three decisive advantages over split inteins from a scientific and therapeutic perspective: i) It shows higher flexibility in the split site selection. ii) The efficiency of reconstitution is less dependent on the position of the split site because it occurs at the transcript level and therefore does not require proper protein folding prior to reconstitution. iii) It does not generate bacterial proteins, reducing the risk of immune responses in treated individuals. Expression of bacterial proteins such as Cas in CRISPR-based approaches may already trigger immune responses in treated individuals^14, 47^. The presence of additional bacterial proteins (e.g., inteins) could result in additive or even synergistic effects and thus increase the risk of overshooting immune responses.

Among the various CRISPR-Cas techniques, CRISPRa is particularly attractive for therapeutic applications as it is mutation-independent and causes very little off-target effects^48–50^. This strategy was successfully applied in several preclinical studies to treat acquired or genetic disorders such as obesity, acute kidney injury, diabetes mellitus, muscular atrophy, retinitis pigmentosa, or epilepsy^15, 48–51^. One of the most powerful CRISPRa modules is Cas9-VPR^28^, which is too large to fit into a single AAV vector. REVeRT is capable to functionally reconstitute full-length Cas9-VPR in various therapeutically relevant tissues and organs (i.e., retina, heart, lung, liver, skeletal muscle and brain) and by different routes of application. Cas9-VPR enables the activation of genes which require high expression to achieve a therapeutic effect and is therefore superior over other less efficient CRISPRa modules (e.g. (Sa)Cas9-VP64^52, 53^) which are small enough to be delivered with a single AAV strategy^49, 50^. In addition, we provide a proof-of-principle that REVeRT can be used efficiently for *in vivo* multiplexing CRISPR-Cas-based approaches aiming at concurrent knockout and activation of different genes. Such approaches enable the treatment of more complex genetic or acquired diseases. These include genetic disorders caused by gain-of-function mutations as demonstrated in this study. Moreover, the CONNACT strategy could also provide a novel platform for treatment of other disorders involving multiple genes or pathways such as cancer or neurodegenerative diseases. For example, REVeRT in combination with CONNACT could be used to knockdown oncogenic genes with simultaneous activation of tumor suppressor genes, or to activate neuroprotective genes with knockdown of cell-damaging or proapoptotic genes.

As REVeRT functions efficiently in different tissues or organs even after intraperitoneal application, it could allow for a more convenient treatment of affected individuals for a given disease, e.g., if local administration routes are inappropriate or if multiple organs have to be transduced simultaneously. Nevertheless, although yielding highly promising results in terms of efficacy, further safety parameters such as tolerability, toxicity or immunoreactivity need to be investigated in non-human primates before REVeRT can be used in first clinical trials. Finally, we show that REVeRT works efficiently after intravitreal injections in the context of classical gene supplementation designed to treat one of the most common genetic blinding diseases caused by mutations in the large *ABCA4* gene. In addition to *ABCA4*, there are many other genes that lead to frequent genetic diseases but exceed the capacity of AAVs. These include genes associated with other diseases of the retina (e.g., Usher syndrome or congenital stationary night blindness), the musculature (e.g., Duchenne muscular atrophy), the hematopoietic system (e.g., hemophilia A), the heart (e.g., catecholaminergic polymorphic ventricular tachycardia), but also multisystem disorders such as Timothy syndrome.

## Methods

### Mice

Animal procedures were performed with permission from local authorities (District Government of Upper Bavaria, Germany) in accordance with the German laws on animal welfare (Tierschutzgesetz). P0 pups were euthanized by decapitation for hippocampal primary neuron preparation. Adult mice were anesthetized with isoflurane and euthanized by cervical dislocation for organ harvest or decapitation for hippocampi isolation. C57BL/6J WT mice, *Rho^+/P23H^* (B6.129S6(Cg)-*Rho^tm1.1Kpal^*/J, Jackson Laboratory, RRID:IMSR_JAX:017628) mice or *Abca4^-/-^ Rdh8^-/-^* (B6;129-*Abca4^tm1Ght^ Rdh8^tm1Kpal^*/J, Jackson Laboratory, RRID:IMSR_JAX:030503) mice homozygous for the Leu450 polymorphism in *Rpe65* were used as indicated.

### sgRNA/pegRNA design

sgRNAs were designed as described previously^15^. pegRNAs were designed using the pegFinder (http://pegfinder.sidichenlab.org/54) for targeting *Rho* exon 1, *Ush2a* exon 12 and *Cnga1* exon 9. For *Dnmt1* exon 1 and the human intergenic *HEK3* locus, previously published pegRNAs were used^32^. sgRNA and pegRNA sequences are shown in Table S4 and Table S5.

### Cloning

The pAAV.CMV.Luc.IRES.EGFP.SV40, dCas9-VPR, pSMVP-Cas9N, pSMVP-Cas9C, pAAV-CMV-Cas9C-VPR, and pCMV-PE2 plasmids were obtained from Addgene (#105533, #63798, #80934, #80939, and #80933, #132775). REVeRT-based reporter genes, (d)Cas9-VPR, PE2 and ABCA4 constructs were synthesized (BioCat GmbH) and cloned into an ITR-containing plasmid (pAAV2.1). The luciferase constructs were generated using the pAAV.CMV.Luc.IRES.EGFP.SV40 plasmid and cloned into pAAV2.1. Binding domains and splice sites were exchanged, promoter and pA signal removed and sgRNA expression cassettes added^31^ using standard cloning techniques. All constructs were sequenced before use (Eurofins Genomics).

### Prediction of splice site strength

NNSplice (Berkeley Drosophila Genome Project, http://www.fruitfly.org/seq_tools/splice.html) and the alternative splice site predictor (ASSP, http://wangcomputing.com/assp/) were used. All used splice sites received high splice site prediction scores (0.91-1.0 for NNSplice and 9.2-4.4 for ASSP).

### Cell culture and transfection

HEK293 cells (DMSZ) and HEK293T cells (Takara) were maintained in Dulbecco’s modified Eagle’s medium (DMEM) GlutaMAX^TM^ (Thermo Fisher Scientific, high glucose for HEK293T) supplemented with 10 % FBS (Biochrom) and with 1 % penicillin/streptomycin (P/S, Biochrom) at 37 °C, 10 % CO_2_. The murine retinoblastoma-derived 661W cell line was provided by M. Al-Ubaidi^55^ and cultured in DMEM GlutaMAX^TM^ medium (with pyruvate) supplemented with 10 % FBS and 1 % Anti-Anti (Thermo Fisher Scientific) at 37 °C, 5 % CO_2_. Immortalized MEF cells were generated as described previously^56, 57^ and cultured in DMEM GlutaMAX^TM^ medium supplemented with 10 % FBS and 1% P/S at 37 °C, 5 % CO_2_. HEK293, HEK293T cells were transfected using the standard calcium phosphate technique. 661W cells were transfected with the calcium phosphate technique or the Xfect^TM^ transfection reagent (Takara). MEF cells were transfected using TurboFect^TM^ Transfection Reagent (Thermo Fisher Scientific) or the Xfect^TM^ transfection reagent. For co-transfections, equimolar amounts of each plasmid were used. Cells were harvested 48 h post transfection.

### Primary neuron preparation and transduction

The preparation of the hippocampus and isolation of primary hippocampal neurons was done as described previously^58^. The primary neuron culture was maintained at 37 °C and 5 % CO_2_ with fresh medium provided every 2-3 days. Cultivation medium consists of 1 % P/S, 1 % GlutaMAX^TM^-I and 20 % B27 Supplement in Neurobasal A medium. Primary neurons were transduced at seven days *in vitro* by adding 7.5 x 10^10^ vg of titer-matched REVeRT AAV8(Y733F) vectors to the media. Neurons were harvested seven days post-transduction in cell lysis buffer (RLT+ buffer (Qiagen) + 1 % β-mercaptoethanol (Merck)).

### AAV production and subretinal/intravitreal injections

REVeRT dual AAV vectors were produced using either the AAV8(Y733F)^27^, AAV9 or AAV2.GL capsid^43^ as described previously^31^. For subretinal and intravitreal injections, WT, *Rho*^P23H/+^ and *Abca4*^-/-^*Rdh8*^-/-^ mice were anaesthetized by intraperitoneal injection of 0.02 mg/g body weight ketamine and 0.04 mg/g body weight xylazine and the pupils were dilated using 1 % atropine and 0.5 % tropicamide eyedrops. Subretinal injections were performed with a NANOFIL 10 µl syringe (World Precision Instruments) and a 34 G beveled needle (World Precision Instruments). For the reconstitution of fluorophores, WT mice were subretinally injected at P21 with 1 µl of dual titer-matched AAV8(Y733F)-split-fluorophore (1.2 x 10^12^ vg/µl) vectors. For transactivation of *Myo7b*, WT mice were subretinally injected at P28-P56 with 1 µl of dual titer-matched AAV8(Y733F)-split-dCas9-VPR (4.71 x 10^10^ vg/µl) vectors. For multiplexed gene knockout and transactivation, WT and *Rho*^P23H/+^ mice were subretinally injected at P21 with 1 µl of dual titer-matched AAV8(Y733F)-split-Cas9-VPR (2.6 x 10^11^ vg/µl) vectors. Contralateral eyes were control injected with 0.9 % NaCl (saline) or phosphate buffer saline containing magnesium and potassium + 0.014% Tween (AAV buffer). To control for Cas9-VPR-specific off-target effects in *Rho*^P23H/+^ mice, AAVs expressing a *lacZ*-targeting sgRNA were injected into contralateral eyes. Eyes were harvested 4 weeks post injection. *Abca4*^-/-^*Rdh8*^-/-^ were injected intravitreally at P14 with 1 µl of dual titer-matched AAV2.GL-split-ABCA4 (1.14 x 10^10^ vg/µl). The contralateral eyes were control injected with AAV buffer. Retinas were isolated after final *in vivo* experiments at 10 weeks post injection.

### Stereotactic injections

Anesthetized WT mice (P30) were fixed in the stereotactic apparatus in sternal recumbence and the skull was uncovered for bregma and lambda identification. Using the Neurostar software (Neurostar GmbH) bregma and lambda were used as reference for calibration. The coordinates for the bilateral injection into the hippocampus were ML ± 1.4, AP −2.06, and DV 1.4. Titer-matched REVeRT AAV8(Y733F) vectors (4.71 x 10^10^ vg in 0.8 µl) were injected into the hippocampus at a flow-rate of 0.3 µl/min. Hippocampi were harvested 4 weeks post injection.

### Intraperitoneal injections

WT mice were injected intraperitoneally at P7 using a 10 µl syringe (Hamilton) and a 28 G beveled needle (Hamilton). For the reconstitution of luciferase, 10 µl of dual titer-matched AAV9 (2 x 10^11^ vg/µl) vectors were injected. For the reconstitution of dCas9-VPR, 10 µl of dual titer-matched AAV8(Y733F) (8 x 10^10^ vg/µl) vectors were injected.

### Luciferase experiments in vitro and in mice

HEK293T cells were transfected as described above. After 72 hours the medium was replaced with 1 ml of culture medium containing 150 mg/ml luciferin. After a 10 min incubation, the luminescence was measured on a bioluminescence imaging system (IVIS Spectrum, PerkinElmer). Luminescence images were taken with automatic exposure time, medium binning and F/stop=1 (Living Image® 4.7.4, PerkinElmer). Three weeks after intraperitoneal injection, 150 mg/kg body weight D-Luciferin (PerkinElmer) were intraperitoneally injected 10-15 minutes prior to imaging. For imaging with the IVIS Spectrum bioluminescence system, the mice were anesthetized with 5 % v/v isoflurane in carbogen at 4 L/min. During the measurement, general anesthesia was maintained by mask inhalation of 1.5 % v/v isoflurane. Luminescence images were taken with automatic exposure time, medium binning and F/stop=1 (Living Image® 4.7.4). After imaging, the mice were sacrificed and organs were dissected. Using the luciferase assay system (Promega), the organ specific luminescence was quantified. Total protein concentration was determined using the Qubit™ Protein and Protein Broad Range (BR) Assay Kit (Invitrogen).

### Luciferase experiments in retinal organoids

Human retinal organoids (hROs) were obtained from Newcells Biotech and maintained in the provided culture medium in a round bottom 96-well plate in a CO_2_ incubator at 37 °C, 5 % CO_2_, 20 % O_2_. The media was renewed every 3-4 days. For transduction, the culture medium was replaced with 50 µl of medium containing 1.5 x 10^12^ vg of a single AAV or 3 x 10^12^ vg of titer-matched dual REVeRT AAV9 vectors expressing the split luciferase. After 7 hours, 150 µl medium were added. The medium was changed every 2-3 days thereafter. Seven days after transduction, the medium was replaced with 100 µl of culture medium containing 150 mg/ml luciferin. After 10 min incubation, the luminescence was measured on IVIS Spectrum bioluminescence system. Luminescence images were taken with automatic exposure time, medium binning and F/stop=1.

### RNA isolation and cDNA synthesis

For RNA extraction from transiently transfected cells, cells were harvested in cell lysis buffer. The retinas were collected as described previously^59^. Retinas and dissected organs (hippocampi, heart, liver, lung, skeletal muscle and whole brain) were shock-frozen in liquid nitrogen and homogenized in cell lysis buffer using the mixer mill MM400 (Retsch). RNA was extracted using the RNeasy Plus Mini Kit (Qiagen) for transfected cells, primary neurons, retinas, hippocampi, heart, liver and lung. For primary neurons, additional on-column DNase I digest was performed. For RNA isolation from skeletal muscle, the RNeasy Fibrous Tissue Mini Kit (Qiagen) was used. For RNA isolation from brain tissue, 25 mg of the homogenized lysate was added to 800 µl TRI-Reagent^TM^ (Zymo Research) and homogenized further using the mixer mill MM400 (Retsch) followed by RNA isolation with the Direct-zol™ DNA/RNA Miniprep Kit (Zymo Research). Total RNA concentrations were measured using the Nanodrop^TM^ 2000c spectrophotometer (Thermo Fisher Scientific). Complementary DNA (cDNA) was synthesized using the RevertAid First Strand cDNA Synthesis Kit (Thermo Fisher Scientific) for up to 1 μg total RNA.

### (q)RT-PCR

For reverse transcription PCR (RT-PCR), the Herculase II polymerase (Agilent Technologies) was used. Quantitative real-time RT-PCR (qRT-PCR) was performed with duplicates on a MicroAmp™ Fast Optical 96-Well Reaction Plate (Themo Fisher Scientific) using the QuantStudio^TM^ 5 Real-Time PCR system (Thermo Fisher Scientific) and the PowerUp^TM^ SYBR^TM^ Green Master Mix (Themo Fisher Scientific). The expression level of all genes was normalized to *Alas* and calculated with the 2^-ΔΔC(T)^ method. All primers are listed in Table S6.

### Protein extraction

For protein extraction from transiently transfected cells, the cells were harvested in TX lysis buffer containing 0.5 % Triton-X (v/v), 3 % NaCl (5 M, v/v), 0.08 % CaCl_2_ (2.5 M, v/v) and cOmplete^TM^ ULTRA Protease Inhibitor Cocktail tablets (Roche) in H_2_O. For the Myo7b western blotting, the eyes were removed, punctured with a 21 G needle at the *ora serrata* and incubated in 4 % paraformaldehyde (PFA) for 5 minutes before removing the cornea, lens and vitreous body under a stereomicroscope. The eyecup was frozen in liquid nitrogen. For the ABCA4 western blotting, retinas were collected as described above. 100 µl of 1x RIPA lysis buffer (Merck) supplemented with 10 % glycerol and a proteasome inhibitor (Roche) was added. Cells, eyecups and retinas were disrupted using the mixer mill MM400. The lysates were rotated end-over-end (VWR^TM^ tube rotator) for 20 min (cerulean, Myo7b) or 60 min (ABCA4) at 4 °C and centrifuged.

### Western blotting

For Western blotting, 15 – 30 μl of the lysates were incubated in 1x Laemmli sample buffer containing DTT at 72°C for 10 min. For ABCA4, 30 µl of lysates were incubated with Protein Loading Buffer 5x (National Diagnostics) at 37°C for 15 min. The proteins were separated on an SDS-polyacrylamide gel via gel electrophoresis. Cerulean and Myo7b blots were stained for 1 h at room temperature (RT) or overnight at 4 °C using the mouse anti-GFP (detects cerulean, 1:2000, JL-8, Clontech, Takara) and the mouse anti-β-tubulin antibody (1:500, D3U1W, Cell Signaling) or with the rabbit anti-Myo7b (1:1000, Sigma-Aldrich) and the mouse anti-β-actin antibody (1:3000, AC-15, Sigma-Aldrich). ABCA4 blots were stained overnight at 4 °C using the rabbit anti-ABCA4 (1:1000, abcam) and the rabbit anti-hexokinase II (1:1000, C64G5, Cell Signaling) antibody. Secondary antibodies (Peroxidase AffiniPure anti-rabbit or anti-mouse IgG, 1:2000, Jackson) were added for 1 h at RT. The western blots were imaged and the relative band intensities quantified using the ImageLab software (Bio-Rad) or VisionWorks LS Analysis Software (ABCA4, Analytik Jena).

### Flow cytometry

Flow cytometry was conducted on HEK293 cells 48 h post transfection. Cells were detached from the culture plate with TrypLE™ Express (Thermo Fisher Scientific) and centrifuged at 100 x g for 5 min. The pellet was resuspended in 400 µl FACS buffer (2 % FBS, 2 mM EDTA (VWR), 25 mM HEPES (Sigma) in PBS), the cells were separated with a 40 µm strainer (pluriSelect) and flow cytometry was performed on a BD LSR Fortessa^TM^ (BD Bioscience) by the Core Facility Flow Cytometry Biomedical Center (LMU Munich). Dead cells were excluded from analysis as determined by SYTOX Blue Dead Cell Stain (Thermo Fisher Scientific). The used gating strategy is shown in Fig. S8.

### Fluorescence-(FHC) and immunohistochemistry (IHC)

The eyes were removed and placed in 0.1 M PB. The eyeball was punctured at the ora serrata with a 21 G cannula and fixed in 4 % PFA for 5 min. The cornea, lens and vitreous body were removed using a stereomicroscope. The eye cup was fixed in 4 % PFA for 45 min at RT, cryopreserved a in 30 % sucrose solution (w/v) and cryosectioned into 10 µm slices. For FHC, the retina was protected from light during the whole procedure.

For different injections, the retinal sections were stained with different antibodies: mouse anti– rhodopsin 1D4 (1:1500, 1D4, Merck), rabbit anti-M-opsin (1:300, Merck) for detection of native M-Opsin in cones and activated M-Opsin in rods, mouse anti-ABCA4 (1:100, 3F4, Santa Cruz) and a custom-made rabbit anti-CNGB1 antibody (1:5000) as an outer segment marker. Fluorescein isothiocyanate–conjugated anti-lectin from Arachis hypogaea [peanut agglutinin (PNA)] (1:100, Sigma-Aldrich, L7381) served as a marker for cones. After overnight incubation at 4 °C, the following secondary antibodies were incubated for 1.5 h at RT: Cy3 anti-rabbit (1:400, Jackson ImmunoResearch), Cy5 anti-mouse (1:400, Jackson ImmunoResearch) and Alexa Fluor Plus 488 anti-mouse (1:800, Thermo Fisher Scientific). No antibodies were used on FHC sections. For nuclear staining, Hoechst 33342 solution (5 µg/ml, Invitrogen) was incubated for 5 min.

### Microscopy

For FHC and Rho^P23H^ IHC retinal sections, z-stack images were obtained using a Leica TCS SP8 inverted confocal laser scanning microscope and the LASX software (Leica). For fluorophore-specific bleaching, a single cell was zoomed in on and the laser intensity of the citrine- and mCherry-specific channels was increased to 80 %. For ABCA4^-/-^ IHC sections, z-stack images were acquired with a ZEISS AxioObserver7 imaging platform with ApoTome2. Deconvolution and maximum intensity z-projection were performed in the ZEISS Zen Blue 3.3 Software. Images of transfected living cells were obtained using the Leica TCS SP8 spectral confocal laser scanning microscope equipped with a HCX APO 20x/1.00 W objective (Leica). All images were processed further with the ImageJ software (National Institutes of Health).

### DNA isolation

Transfected HEK293T, 661W and MEF cells were harvested in cell lysis buffer. The cells were disrupted using the mixer mill MM400. The lysates were centrifuged at 5000 x g for 10 min at 4 °C and supernatant added to gDNA extraction columns (Zymo Research). The columns were centrifuged for 4 min at 1500 x g and 1 min at 10000 x g at RT and incubated with 400 µl Genomic lysis buffer (Zymo Research) supplemented with RNase A (Sigma-Aldrich, 1:5000) for 10 min followed by a 2 min centrifugation at RT. 400 µl of DNA Pre-Wash buffer (Zymo Research) and 600 µl of g-DNA Wash buffer (Zymo Research) were added with intermediate centrifugation steps for 1 min at 10000 x g. 600 µl of g-DNA Wash buffer was added again and centrifuged for 2 min at 10000 x g. To elute gDNA, 100 µl of H_2_O were added to the column, incubated for 20 min at RT and centrifuged for 2 min at 10000 x g. gDNA concentrations were measured using the NanodropTM 2000c spectrophotometer.

### Quantification of prime editing efficiency

For amplification of the respective target locus, 100 ng of the extracted gDNA were for PCR with the Q5 high-fidelity DNA polymerase (New England Biolabs). The product was analyzed via gel electrophoresis, purified from the agarose gel with the QIAquick gel extraction kit (Qiagen) and Sanger sequenced (Eurofins Genomics). To quantify the prime editing efficiency, the PCR was repeated using primers containing Illumina partial adapters (Table S6). The product was purified as describe above and sent to next-generation sequencing (Azenta Life Sciences). The editing efficiency was calculated from the total number of reads per sample and the total number of correctly edited reads per sample.

### RNA-Seq

Total RNA was sent to a commercial provider (Azenta Life Sciences) to perform RNA sequencing and analysis. mRNA was enriched by poly(A) selection. Paired-end sequencing was performed on an Illumina HiSeq system with > 20 million reads per sample and 150 bp read length. Trimmomatic v.0.36 was used to trim the sequence reads for adapter sequences and quality. The reads were mapped to the *Mus musculus* GRCm38 reference genome using the STAR aligner v.2.5.2b. Unique gene hit counts were calculated using FeatureCounts from the Subread package v.1.5.2 and only unique reads mapped to exonic regions were included. DESeq2 was used to analyze the obtained gene hit counts. The Wald test was used to obtain p-values and log2 fold changes. Differentially expressed genes were defined by an absolute log2 fold change > 1 and adjusted p-value < 0.05.

### Electroretinography (ERG)

Mice were dark-adapted overnight. Tropicamide eye drops were applied for pupil dilation (Mydriadicum Stulln, Pharma Stulln GmbH). Full-field ERG responses were recorded using a Celeris apparatus (Diagnosys LLC). Light guide electrodes were placed on each eye. For the scotopic measurements, the eyes were illuminated in single-flashes with increasing light intensities (0.003, 0.01, 0.03, 0.1, 0.3, 1, 3, 10 cd.s/m^2^). After a 5 min-light adaptation step with 9 cd/m^2^, sequential photopic responses were recorded for single-flash steps 0.01, 0.03, 0.1, 0.3, 1, 3, 10 cd.s/m^2^ with constant 9 cd/m^2^ background illumination. Flicker recordings were performed for 20 and 30 Hz at 3 cd.s/m^2^ with constant 9 cd/m^2^ background illumination.

### Optical coherence tomography (OCT) and fundus autofluorescence imaging

OCT and fluorescent confocal scanning laser ophthalmoscopy (cSLO) were performed with an adapted Spectralis HRA + OCT system (Heidelberg Engineering) in combination with optic lenses. OCT scans were conducted with a linear scan mode. The mean photoreceptor layer thickness was calculated from single values measured in the dorsal, temporal, nasal and ventral region around the optic nerve. cSLO was performed directly after OCT measurements. Images were recorded by averaging 100 consecutive imaged using the automated real time mode.

### Quantification and statistical analysis

All values are given as mean ± SEM. The number of replicates (n) can be inferred from scatter plots for each experiment. Statistical analysis was performed using GraphPad PRISM (GraphPad Software). Normality was tested using a Shapiro-Wilk test. For parametric data, unpaired t-test with Welch’s correction or Welch ANOVA was used. For non-parametric data, Mann-Whitney or Kruskal-Wallis test was used. Multiple comparisons were corrected for using

Dunnett’s T3 test. Correlation coefficients were calculated using Spearman’s correlation. In experiments with paired samples, a paired t-test with Welch’s correction or a two-way ANOVA with Šídák’s multiple comparisons test were used.

## Data availability

The datasets generated in this study are available in public repositories or are directly included in the published article (and its supplementary information files). The RNA-Seq data have been deposited to GEO (Gene Expression Omnibus, https://www.ncbi.nlm.nih.gov/geo/) under the GEO accession GSE198893. The NGS data from the prime editing experiments have been also deposited to GEO under the GEO accession GSE198863. All final REVeRT sequence elements used in this study are listed in Table S2. Additionally, sgRNAs, pegRNAs and primers used in this study can be found in Table S4, S5 and S6. Additional data related to this paper are available from the corresponding author upon reasonable request.

## Acknowledgements

We thank B. Noack, J. Koch, and K. Skokann for technical support. We thank S. Mir-Bashiri for her contribution to the *in vitro* characterization of REVeRT and M. Al-Ubaidi for the gift of the 661W cells. Moreover, we thank the Core Facility Flow Cytometry at Biomedical Center, LMU, for performing the flow cytometry experiments. This work was funded by the German Research Foundation (Deutsche Forschungsgemeinschaft, DFG), BE 4830/2-2 and BE4830/5-1 (to E.B.), BI 484/7-2 (to M.B.), MI 1238/4-1 (to S.M.), FE 1929/1-1 (to S.F.), by the German Center for Cardiovascular Research (81X3600702 to S.F.) and by the Swiss National Science Foundation (310030E_213089 to E.B.).

## Author contributions

Conceptualization, E.B.; Methodology, L.M.R. and E.B.; Investigation, L.M.R, K.S.H., S.B.T., N.K., S.B., D.M.M., V.J.W., D.O., V.S., M.Br., N.B., and V.M.; Writing – Original Draft, E.B.; Writing – Review & Editing, L.M.R., K.S.H., S.B.T., N.K., V.M., S.M., S.F. and M.Bi.; Visualization, L.M.R., K.S.H. and E.B.; Funding Acquisition, J.W., S.M., S.F., M.Bi. and E.B.; Supervision, J.W., V.M., S.M., S.F., M.B. and E.B.

## Materials & Correspondence

Correspondence and requests for materials should be addressed to Elvir Becirovic or Lisa Riedmayr.

