## Supplementary information for "mRNA trans-splicing dual AAV vectors for (epi)genome editing and gene therapy"

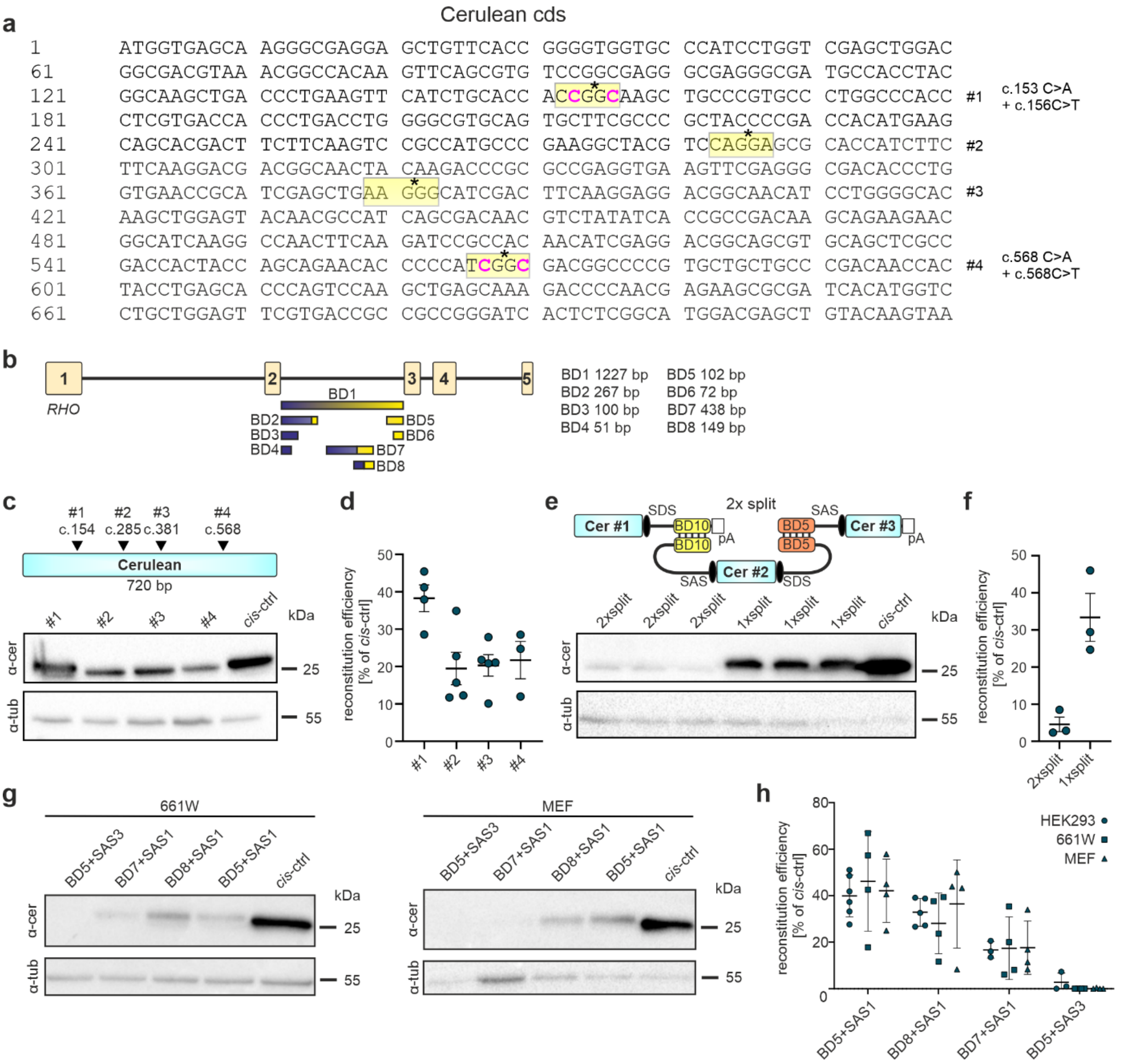


**Fig. S1 *In vitro* evaluation of REVeRT.** **a** Coding sequence (cds) of cerulean. The split sites (#1-4) used are highlighted in yellow. Asterisks mark the exact split position. Introduced silent mutations are highlighted in pink. **b** Scheme depicting *RHO* intron 2 used for BD1-BD8. The respective sequences originate from different regions of the intron as indicated by the color gradient. BD2, BD7 and BD8 represent fusion sequences containing the parts of the 5' end and parts of the 3' end of the intron. **c** Upper panel, Position of the split sites within the cerulean sequence. Lower panel, Western Blot of HEK293 cells transfected with cerulean vectors split at positions #1-4. **d** Ratiometric quantification of reconstitution efficiency relative to the cis-ctrl determined from western blots **c**. **e** Upper panel, Scheme depicting mRNA reconstitution for cerulean split into three fragments. Lower panel, Western Blot of HEK293 cells transfected with cerulean constructs split into two (1xsplit) or three fragments (2xsplit). **f** Ratiometric quantification of reconstitution efficiency relative to the cis-ctrl determined from western blots **e**. **g** Western blot of 661W (left) or mouse embryonic fibroblast (MEF, right) cells transfected with split cerulean constructs in presence of different BDs or SASs. **h** Ratiometric quantification of the reconstitution efficiency relative to the cis-ctrl determined from western blots **g**. The corresponding HEK293 western blots are shown in Fig 1. Scatter plots show mean ± SEM.

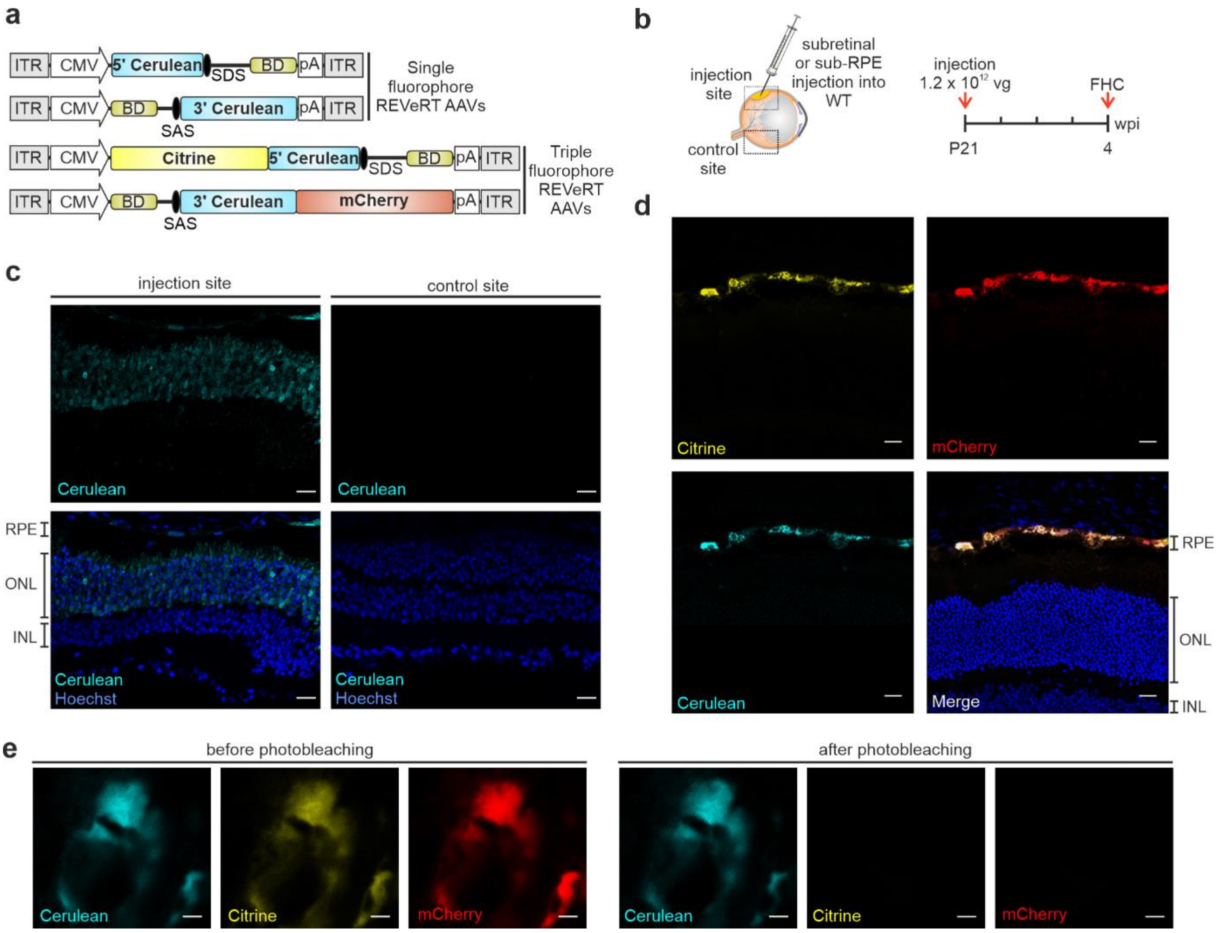


**Fig. S2 Reconstitution of fluorescent reporter genes via dual REVeRT AAVs *in vivo*. a** Design of dual REVeRT AAVs used for experiments shown in **c** (single fluorophore), **d** and **e** (triple fluorophore). ITR, inverted terminal repeats. **b** Experimental setup. FHC, fluorescence histochemistry. wpi, weeks post injection. **c** Representative FHC images of retinas obtained from mice injected with single fluorophore REVeRT AAVs. Left panel, cerulean expression along the injection site; Right panel, sections from the non-injected control site within the same retina. RPE, retinal pigment epithelium; ONL, outer nuclear layer; INL, inner nuclear layer. Scale bar, 20 µm. **d**, **e** FHC of a retina injected with triple fluorophore REVeRT AAVs (citrine, split cerulean and mCherry) system. Scale bar, 30 µm. **e** High magnification images of RPE cell expressing all three fluorophores before and after selective photobleaching of the mCherry and citrine signal. Photobleaching confirmed the presence of reconstituted cerulean. Scale bar, 2 µm. All results in **c-e** are derived from non-amplified (i.e., antibody-free) fluorophore signals.

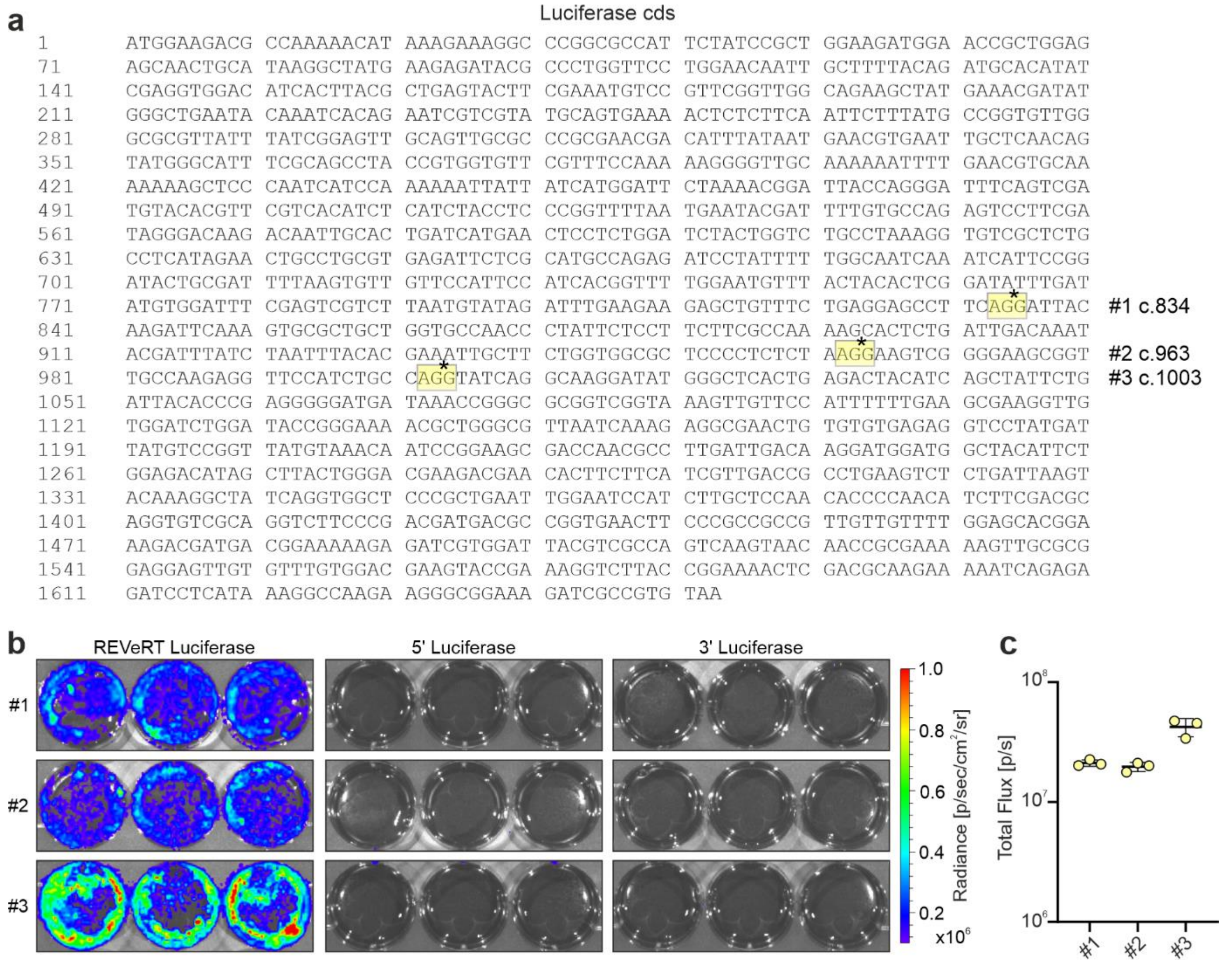


**Fig. S3 Split luciferase reporter assay. a** Luciferase coding sequence (cds). The split sites (#1-#3) used in this study are highlighted in yellow. Asterisks mark the exact split position. **b** Luminescence images of HEK293T cells transfected with luciferase constructs split at different positions as indicated. The cells were either transfected with dual vectors (REVeRT Luciferase) or with a single vector (5’ Luciferase, 3’ Luciferase) as negative control. **c** Quantification of the results shown in **b**. Scatter plot shows mean ± SEM.

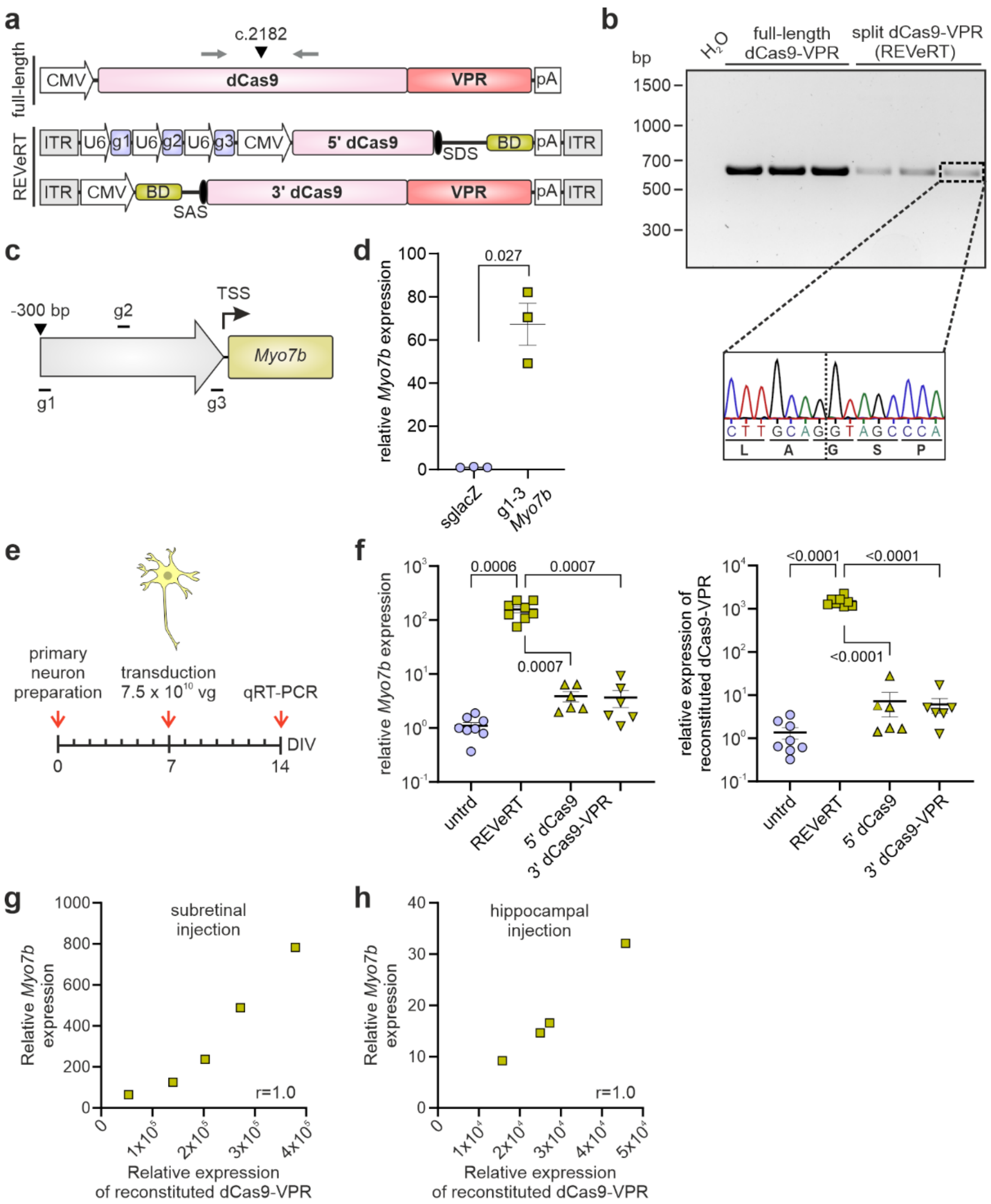


**Fig. S4** ***Myo7b*** **transactivation using split dCas9-VPR reconstituted via REVeRT. a** Split (REVeRT) and full-length dCas9-VPR constructs used in **b**-**h**. Split site (arrowhead) and junction-spanning primers (arrows) are indicated. **b** RT-PCR using indicated primer pair. Sequencing of the splicing junction is shown below. **c** Binding positions of the three sgRNAs targeting the region upstream of the transcription start site (TSS) of the *Myo7b* gene. **d** qRT-PCR from 661W cells co-transfected with split dCas9-VPR and either *lacZ*- or *Myo7b*-targeting (g1-g3) sgRNAs. *lacZ*-transfected cells were used as a reference. A Mann-Whitney test was used for statistical analysis. **e** Time scale for experiments in mouse primary neurons transduced with dual REVeRT AAV8Y733F vectors. DIV, days *in vitro*. **f** Relative expression of *Myo7b* and reconstituted dCas9-VPR in transduced mouse primary neurons. Welch ANOVA with Dunnett T3 was used for statistical analysis. Untreated cells (untrd) were used as a reference. Neurons transduced with single vectors (5' dCas9, 3' dCas9-VPR) served as controls**.** **g**, **h** Dose-dependency between dCas9-VPR reconstitution and transcriptional activation of *Myo7b* upon subretinal (**g**) or hippocampal injection (**h**). Spearman correlation coefficient is in the lower right corner. Scatter plots show mean ± SEM.

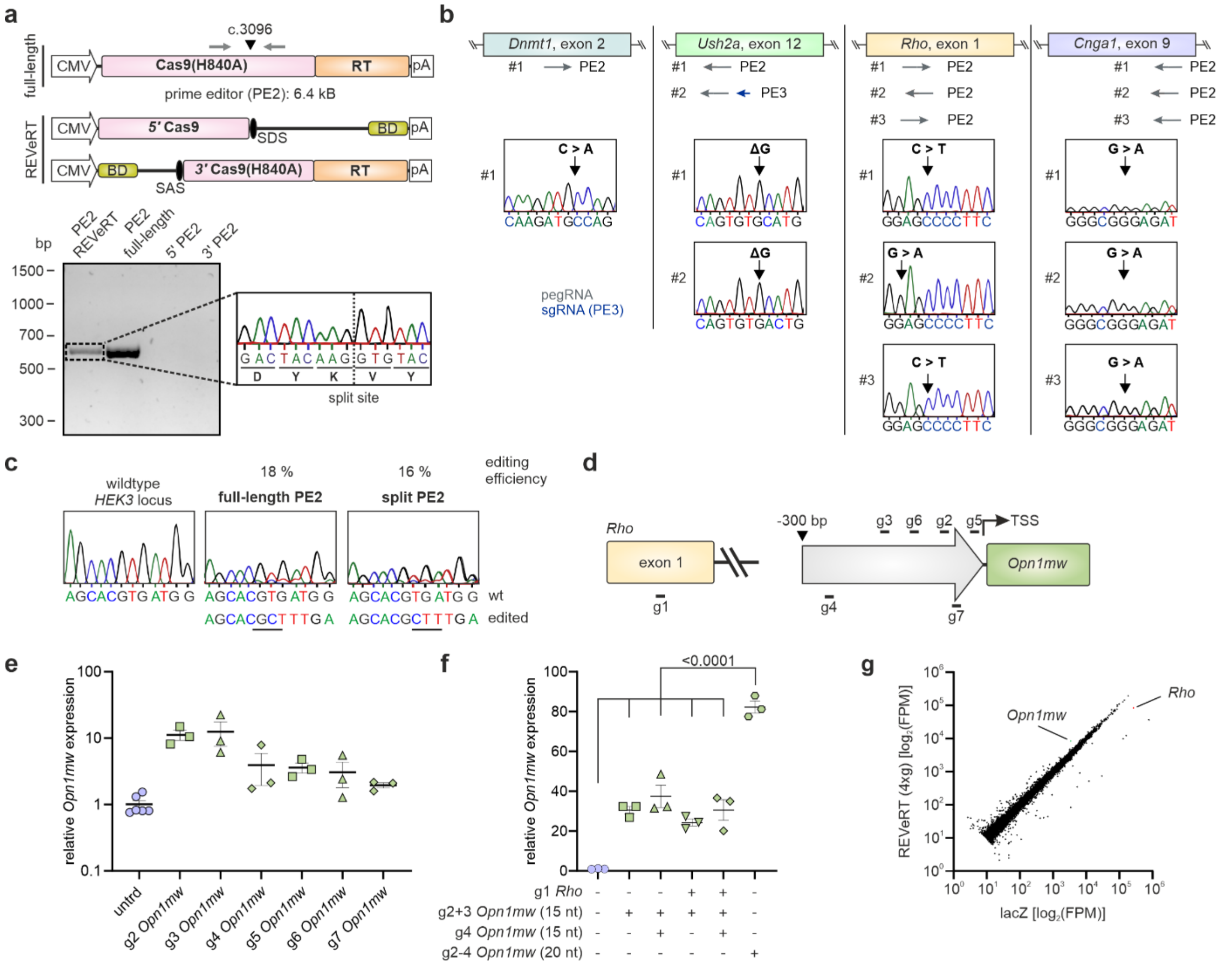

**Fig. S5 *In vitro* prime editing and CONNACT strategy via REVeRT. a** Upper panel, split and full-length prime editor 2 (PE2) constructs used in **b** and **c**. Split site (arrowhead) and junction-spanning primers (arrows) are indicated. Lower panel, RT-PCR of reconstituted PE2 using junction-spanning primers and sequencing result of the splicing junction. Single vectors (5' PE2 and 3' PE2) served as controls. **b** Sequencing results of the targeted genomic loci after co-transfection of 661W and MEF cells with PE2 or PE3 (PE2 + additional sgRNA introducing a nick) and corresponding pegRNAs. Target exons and binding position of the pegRNAs are depicted above. **c** Sequencing of the *HEK3* locus after co-transfection of HEK293T cells with split or full-length PE2 and pegRNA. **d** Binding positions of the sgRNAs targeting *Rho* exon 1 (g1) or the region upstream of the TSS of *Opn1mw* (g2-g7). **e** qRT-PCR from MEF cells co-transfected with dCas9-VPR and g2 – g7. Untreated cells (untrd) served as reference. **f** qRT-PCR from MEF cells co-transfected with dCas9-VPR and different combinations of sgRNAs as indicated. Spacer lengths of *Opn1mw*-targeting sgRNAs are shown in brackets. Untreated cells served as reference. One-way ANOVA with Dunnett's T3 test was used to compare expression levels using a 15 nt vs. 20 nt spacer. Scatter plots show mean ± SEM. **g** RNA-Seq analysis of retinas originating from three different animals (dark pink data points in Fig. 4d) injected with REVeRT(4xg) or *lacZ* vectors. *Opn1mw* and *Rho* transcripts are highlighted. FPM, fragments per million.

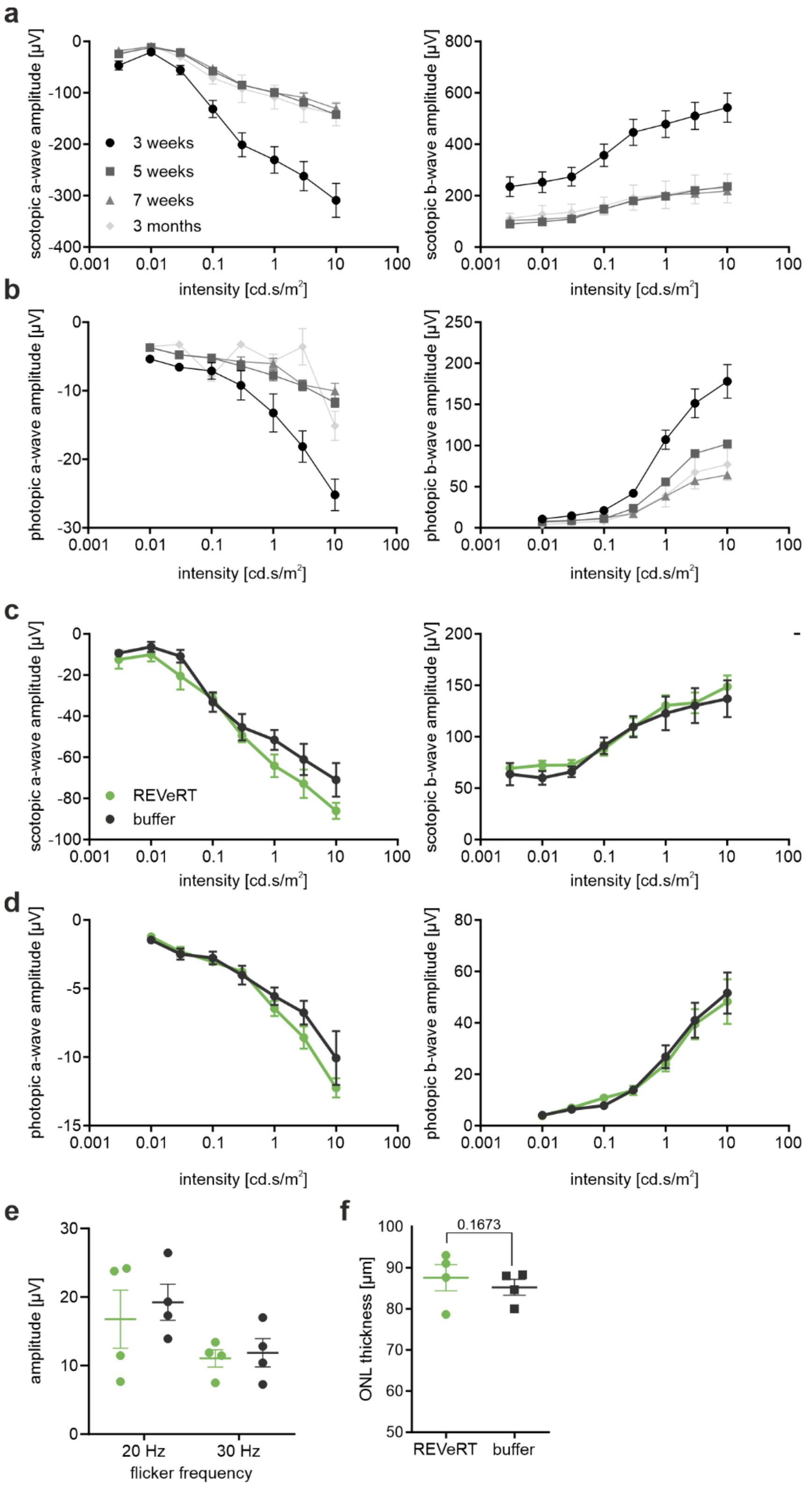


**Fig. S6 Natural history and gene therapy in Abca4-deficient mice. a**, **b** Scotopic (**a**) and photopic (**b**) ERG measurements for untreated *Abca4*$^{-/-}$/*Rdh8*$^{-/-}$ mice between 3 weeks and 3 months of age as indicated. **c**, **d** ERG measurements of injected *Abca4*$^{-/-}$/*Rdh8*$^{-/-}$ mice at 4 wpi under scotopic (**c**) and photopic (**d**) conditions. **e** Amplitudes from flicker stimuli at 20 Hz and 30 Hz under photopic conditions at 4 wpi. **f** OCT measurements of injected *Abca4*$^{-/-}$/*Rdh8*$^{-/-}$ mice at 4 wpi. A two-tailed paired t-test was used for statistical analysis. Plots show mean ± SEM.

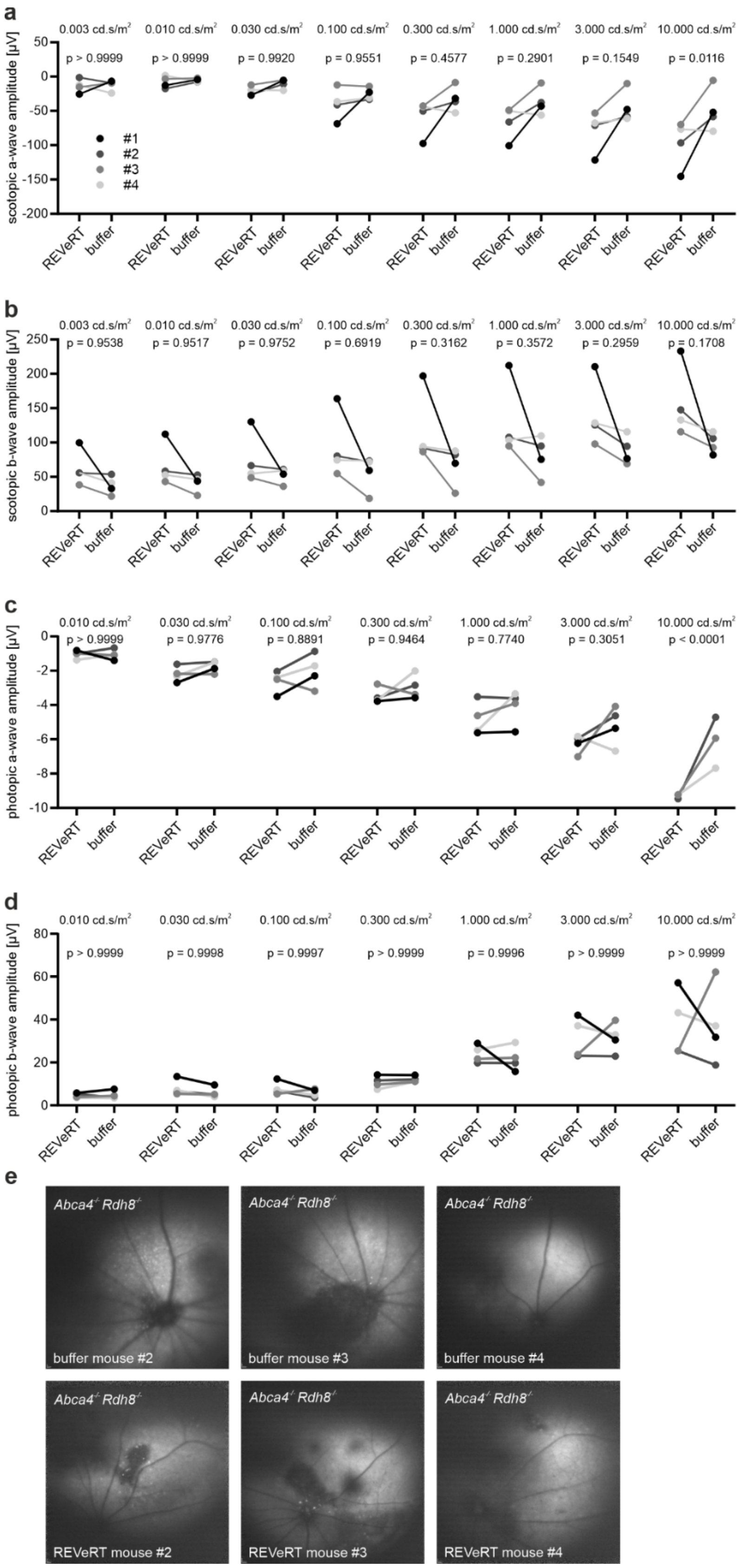


**Fig. S7 Pairwise comparison of treated and buffer-injected eyes in Abca4-deficient mice.** **a**-**d** Pairwise plots for the ERG measurements under scotopic (**a**, **b**) or photopic (**c**, **d**) conditions as indicated. Light intensities and p-values (two-way ANOVA with Šídák's multiple comparison test) are shown above the corresponding measurements. **e** SLO images of mice #2-4 at 10 wpi.

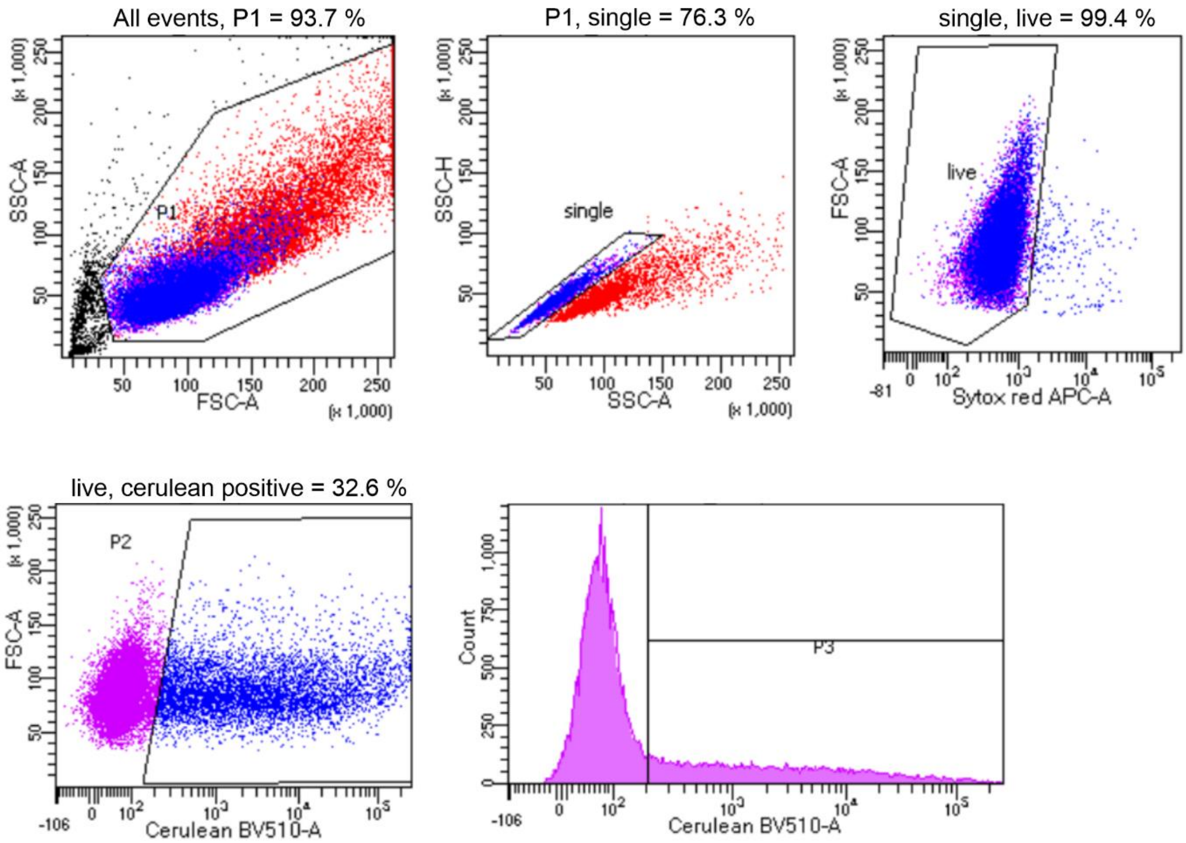


**Fig. S8 Gating Strategy used for flow cytometry on HEK293 cells transfected with split cerulean constructs.**

**Table S1. Binding domain analysis.**

| Binding domain | Reconstitution efficiency WB ± SEM [%] | Reconstitution efficiency FC ± SEM [%] | Length [bp] | GC content [%] |
|---|---|---|---|---|
| 1 | 2.1 ± 2.1 | 1.7 ± 0.9 | 1227 | 55.3 |
| 2 | 27.0 ± 5.8 | 17.3 ± 1.2 | 267 | 55.1 |
| 3 | 5.7 ± 1.8 | 4.2 ± 0.4 | 100 | 57.0 |
| 4 | 37.3 ± 5.4 | 31.5 ± 1.7 | 51 | 56.9 |
| 5 | 39.9 ± 2.3 | 36.4 ± 1.4 | 102 | 62.7 |
| 6 | 24.2 ± 6.6 | 12.1 ± 0.2 | 72 | 56.9 |
| 7 | 16.7 ± 2.1 | 13.6 ± 0.5 | 438 | 56.8 |
| 8 | 32.9 ± 2.5 | 35.8 ± 1.8 | 105 | 59.0 |
| 9 | 29.2 ± 9.4 | 24.0 ± 0.7 | 100 | 47.0 |
| 10 | 53.5 ± 5.7 | 44.0 ± 1.3 | 100 | 46.0 |
| 11 | 30.7 ± 5.0 | 20.3 ± 0.7 | 100 | 48.0 |
| 12 | 13.2 ± 5.0 | 16.8 ± 0.5 | 100 | 48.0 |

**Table S2. Trans-splicing elements used for REVeRT.**

| Element | Sequence |
|---|---|
| BD: BD10 | CATCTGACCACC**T**GCGAA(Δ5bp)TTTTTGCATC**CT**GCTG**TT**TAAT **CT**GCGTTGG(Δ3bp)TTTAACCGCC**T**GTC**T**GGCTTT**T**TTTCAC**T**GA TGTG(Δ1bp)AT*CGCCTTGATGCACT* |
| SAS: SAS1 | CAACGAGTCTTTTGTCATCTACAG\|G |
| SDS: SDS1 | AG\|GTAAG |

BD domain contains following modifications compared to the native lacZ sequence: 11x substitution (bold), 3x deletion (Δ), 1x artificial sequence (*italic*).

**Table S3. Differentially expressed genes.**

| Gene | ID | log2FoldChange | pvalue | padj |
|---|---|---|---|---|
| *Rad54b* | ENSMUSG00000078773 | 2.36 | 1.72E-16 | 3.79E-13 |
| *Manba* | ENSMUSG00000028164 | 1.77 | 8.14E-38 | 1.08E-33 |
| *Slc6a2* | ENSMUSG00000055368 | 1.65 | 1.88E-29 | 1.24E-25 |
| *Gm19410* | ENSMUSG00000109372 | 1.59 | 4.31E-10 | 5.18E-07 |
| *Frrs1* | ENSMUSG00000033386 | 1.46 | 5.76E-13 | 1.09E-09 |
| *Podn* | ENSMUSG00000028600 | 1.39 | 6.49E-12 | 1.07E-08 |
| ***Opn1mw*** | ENSMUSG00000031394 | 1.29 | 1.87E-25 | 8.22E-22 |
| *Ifi47* | ENSMUSG00000078920 | 1.20 | 1.09E-06 | 5.61E-04 |
| *Gm16062* | ENSMUSG00000087249 | 1.18 | 6.95E-07 | 4.48E-04 |
| *Iigp1* | ENSMUSG00000054072 | 1.13 | 1.91E-06 | 7.88E-04 |
| *BC065403* | ENSMUSG00000097211 | 1.13 | 1.40E-04 | 1.01E-02 |
| *Cutal* | ENSMUSG00000026870 | 1.09 | 6.47E-06 | 1.68E-03 |
| *1500026H17Rik* | ENSMUSG00000097383 | 1.09 | 9.25E-05 | 7.69E-03 |
| *Fam204a* | ENSMUSG00000057858 | 1.09 | 6.62E-17 | 1.75E-13 |

| | | | | |
|---|---|---|---|---|
| *Shisa2* | ENSMUSG00000044461 | 1.07 | 4.18E-18 | 1.38E-14 |
| *Gucy1a2* | ENSMUSG00000041624 | -1.07 | 1.00E-03 | 2.95E-02 |
| *Malat1* | ENSMUSG00000092341 | -1.10 | 1.77E-03 | 3.98E-02 |
| *Klf12* | ENSMUSG00000072294 | -1.17 | 2.02E-06 | 7.97E-04 |
| *Kcnh5* | ENSMUSG00000034402 | -1.18 | 3.85E-04 | 1.72E-02 |
| *Dok6* | ENSMUSG00000073514 | -1.28 | 5.92E-05 | 5.79E-03 |
| *Gm11808* | ENSMUSG00000068240 | -1.28 | 7.70E-04 | 2.51E-02 |
| *Gm340* | ENSMUSG00000090673 | -1.55 | 6.85E-06 | 1.68E-03 |
| ***Rho*** | ENSMUSG00000030324 | -1.69 | 6.85E-06 | 1.68E-03 |
| *Cdr1* | ENSMUSG00000090546 | -2.22 | 4.37E-04 | 1.84E-02 |
| *Gm22009* | ENSMUSG00000089417 | -2.29 | 1.11E-04 | 8.76E-03 |
| *Gm26917* | ENSMUSG00000097971 | -2.81 | 3.22E-04 | 1.58E-02 |
| *Scarna2* | ENSMUSG00000088185 | -3.37 | 6.60E-05 | 6.23E-03 |
| *Hist1h4d* | ENSMUSG00000061482 | -3.59 | 5.06E-04 | 2.00E-02 |
| *Rmrp* | ENSMUSG00000088088 | -3.68 | 1.87E-03 | 4.05E-02 |
| *Gm23935* | ENSMUSG00000076258 | -3.71 | 3.06E-04 | 1.51E-02 |
| *Lars2* | ENSMUSG00000035202 | -3.72 | 2.85E-04 | 1.45E-02 |
| *Rn7sk* | ENSMUSG00000065037 | -3.94 | 4.07E-04 | 1.76E-02 |
| *Gm24187* | ENSMUSG00000088609 | -3.96 | 1.35E-04 | 1.00E-02 |
| *Gm42418* | ENSMUSG00000098178 | -4.42 | 3.62E-09 | 3.42E-06 |
| *Gm24270* | ENSMUSG00000076281 | -4.55 | 4.02E-05 | 4.62E-03 |
| *Mir6236* | ENSMUSG00000098973 | -5.32 | 1.12E-05 | 2.13E-03 |
| *Gm15564* | ENSMUSG00000086324 | -5.37 | 6.06E-06 | 1.61E-03 |

Significant up- or downregulation is defined as absolute log2 fold change > 1 and adjusted p-value < 0.05.

**Table S4 sgRNA sequences**

| sgRNA | Spacer sequence | Target and purpose | Figure |
|---|---|---|---|
| g1 Myo7b | AGACTCCAAGAACGCCAGTC | Transcriptional activation of *Myo7b* | 3, S4 |
| g2 Myo7b | GGGCACCATTAACCACTGCT | Transcriptional activation of *Myo7b* | 3, S4 |
| g3 Myo7b | GGAAGGGCTCCAAGCGGAAC | Transcriptional activation of *Myo7b* | 3, S4 |
| g1 Rho | GTACGGTGACGTAGAGCGTG | *Rho* exon 1, Knockout of *Rho* | 4, S5 |
| g2 Opn1mw | GGGGCCTTTAAGGTA | Transcriptional activation of *Opn1mw* | 4, S5 |
| g3 Opn1mw | GCCACCCCTGTGGAT | Transcriptional activation of *Opn1mw* | 4, S5 |
| g4 Opn1mw | CTTGCTTGTTTACAA | Transcriptional activation of *Opn1mw* | 4, S5 |
| g5 Opn1mw | GTCCTGTAACCCCAT | Transcriptional activation of *Opn1mw* | S5 |

| | | | |
|---|---|---|---|
| g6 Opn1mw | GATGATCTAAGTCCT | Transcriptional activation of *Opn1mw* | S5 |
| g7 Opn1mw | CTGCAGGATCAGCCC | Transcriptional activation of *Opn1mw* | S5 |
| *Ush2a* exon12 #2 | CAGACCTCACAAGCACTCCA | *Ush2a* exon 12, Nicking gRNA for PE3 | S5 |

**Table S5 pegRNA sequences used in Fig. S5.**

| pegRNA | Spacer sequence | 3' extension | RT template length (nt) | PBS length (nt) |
|---|---|---|---|---|
| *HEK3*_CTT | GGCCCAGACTGA GCACGTGA | TCTGCCATCAAAGC GTGCTCAGTCTG | 13 | 13 |
| *Dnmt1*_5d_5 GtoT | CGGGCTGGAGC TGTTCGCGC | AAGATGCAAGCGC GAACAGCTCCAG | 12 | 13 |
| *Ush2a* exon 12 #1 + #2 | AACTCTGTGATC CGCTTTCT | TGACACTGCCCAGA AAGCGGATCACAGA | 15 | 13 |
| *Rho* exon 1 #1 | AGTACTGCGGCT GCTCGAAG | GGAGTCCCTTCGAG CAGCCGCAG | 10 | 13 |
| *Rho* exon 1 #2 | CCAACGTCACAG GCGTGGTG | AGGGGCTTCGCAC CACGCCTGTGACG | 13 | 13 |
| *Rho* exon 1 #3 | AGTACTGCGGCT GCTCGAAG | GTGCGGAGTCCCTT CGAGCAGCCGCA | 14 | 12 |
| *Cnga1* exon 9 #1 | CAAGAAAGGGGA CATCGGGC | GTACATCTCTCGCC CGATGTCCCCTTT | 14 | 13 |
| *Cnga1* exon 9 #2 | TATGCAAGAAAG GGGACATC | TACATCTCCCGTCC GATGTCCCCTTTC | 17 | 10 |
| *Cnga1* exon 9 #3 | CAAGAAAGGGGA CATCGGGC | GTACATCTCTCGCC CGATGTCCCCTT | 14 | 12 |

**Table S6 Primer sequences**

| Primer | Sequence (5' – 3') |
|---|---|
| Cas9 forward | AGAACGCTTGAAAACTTACGCT |
| Cas9 reverse | TTGATGTCCAGTTCCTGATCC |
| Cerulean forward | ATGGTGAGCAAGGGCGAGG |
| Cerulean reverse | CTTGTACAGCTCGTCCATGCC |
| Cnga1 forward | TGTGAAGCTGGTCTGTTGGT |
| Cnga1 reverse | CCTCCCTTTCTCTTCCAGCA |
| Dnmt1 forward | GTGTGGTACATGCTGCTTCCG |
| Dnmt reverse | CTCCCTCAAGCTCCCAGTCAAT |
| HEK3 forward | GCATGCATTTGTAGGCTTGA |
| HEK3 reverse | CTTTTCCTCTGTTGAGCTCG |
| HEK3 + adapter forward | ACACTCTTTCCCTACACGACGCTCTTCCGATCTGC ATGCATTTGTAGGCTTGA |

| | |
|---|---|
| HEK3 + adapter reverse | GACTGGAGTTCAGACGTGTGCTCTTCCGATCTCT TTTCCTCTGTTGAGCTCG |
| ITR2 forward | GGAACCCCTAGTGATGGAGTT |
| ITR2 reverse | CGGCCTCAGTGAGCGA |
| PE2 forward | CGATGTGGACGCTATCGTGC |
| PE2 reverse | GCCTTGCCGATTTCCTGCTC |
| qRT-PCR Alas forward | TCGCCGATGCCCATTCTTATC |
| qRT-PCR Alas reverse | GGCCCCAACTTCCATCATCT |
| qRT-PCR Myo7b forward | GGGACACAAGTACAGGAAGGA |
| qRT-PCR Myo7b reverse | GCGTTCAAAGCCCACTAGG |
| qRT-PCR Cas9 reconst forward | AGCACAAGTTTCTGGCCAGGG |
| qRT-PCR Cas9 reconst reverse | GGTAGTTTGGTTCTCTCGGGCC |
| qRT-PCR Opn1mw forward | GGAGCAGGTACTGGCCTTATG |
| qRT-PCR Opn1mw reverse | GGAGGTAGCAGAGCACGATG |
| qRT-PCR Rho forward | GGATCATGGCGTTGGCCTGT |
| qRT-PCR Rho reverse | CCGCATGAACATTGCATGCCC |
| Rho forward | AGCCTTGGTCTCTGTCTACG |
| Rho reverse | TGGTGAATCCTCCGAAGACC |
| Ush2a forward | CCAGGGCTTAGATGCAATCG |
| Ush2a reverse | AAGTAGCCTGCCTTACACTG |
